# An inverse modelling study on the local volume changes during early morphoelastic growth of the fetal human brain

**DOI:** 10.1101/2020.10.08.332411

**Authors:** Z. Wang, B. Martin, J. Weickenmeier, K. Garikipati

## Abstract

We take a data-driven approach to deducing the local volume changes accompanying early development of the fetal human brain. Our approach uses fetal brain atlas MRI data for the geometric changes in representative cases. Using a nonlinear continuum mechanics model of morphoelastic growth, we invert the deformation obtained from MRI registration to arrive at a field for the growth deformation gradient tensor. Our field inversion uses a combination of direct and adjoint methods for computing gradients of the objective function while constraining the optimization by the physics of morphoelastic growth. We thus infer a growth deformation gradient field that obeys the laws of morphoelastic growth. The errors between the MRI data and the forward displacement solution driven by the inverted growth deformation gradient field are found to be smaller than the reference displacement by well over an order of magnitude, and can be driven even lower. The results thus reproduce the three-dimensional growth during the early development of the fetal brain with controllable error. Our findings confirm that early growth is dominated by in plane cortical expansion rather than thickness increase.

## 1 Introduction

Like other organs, the fetal human brain undergoes large changes in volume and geometry during development *in utero*. A foundational understanding of these growth-induced changes can be gained from a *morphoelastic* treatment. Such an approach underlies the now accepted model of morphological development of most biological structures: Mass accretes, either due to cell growth and division, or from the deposition of extra-cellular matrix elements. Due to the elasticity of the newly grown (accreted) tissue, some energy is stored in it, and the relaxation of this energy occurs via an expansion of the tissue. The brain’s grey and white matter are soft materials with molecular structures that are subjected to stress-dependent breakage of secondary bonds, and furthermore, are fluid-filled. There is, therefore, a rate-dependence to the mechanical response of the brain’s constituent matter. However, on the time scales of days to weeks over which the brain undergoes morphological changes, viscous effects are fully relaxed, and elasticity prevails. Specifically, hyperelastic models governed by the equations of nonlinear elasticity describe the mechanical changes accompanying growth.

This is the foundation for the morphoelastic theory of growth, which relies upon a *growth deformation tensor* as one component of a multiplicative decomposition of the total deformation gradient tensor. In general, it is incompatible, meaning that it cannot be expressed as the gradient of a smooth vector field. However, the product obtained by pre-multiplying it with the elastic deformation gradient tensor is indeed compatible, since it expresses the total deformation gradient. The morphoelastic theory of growth has gained interest over the last two decades from the standpoint of neurodevelopmental studies that seek to explain the folding of the brain.

Folding, or sulcification and gyrification, of the brain is common in mammals including primates, cetaceans, pachyderms and ungulates. Folds form in the cortical layer of grey matter, and in species such as humans that demonstrate pronounced gyrencephaly, the sulci can be significantly deeper than the cortical thickness. A folded cortex confers a cognitive advantage by increasing the surface area enclosed within the skull, translating to greater capacity for intelligence. Normally developed human brains have a gyrification index (ratio of actual surface area to the surface area of an enveloping surface) approaching 2.55 [47]. Neurodevelopmental pathologies are associated with significant departures from this value. In humans, polymicrogyria (shallow, more frequent folding) is associated with developmental delays and epilepsy [24]. Pachygyria (shallow, less frequent and flatter folds) is associated with seizures, cognitive impairment and in rare cases, afflictions such as bipolar disorder [36]. Lissencephaly (abscence of folds) is associated with abnormal EEG patterns, intractable epilepsy [25] and cognitive impairment [29].

Fetal MRI data indicate that the human brain is almost perfectly smooth until 24 weeks of gestation [22, 16, 18], from which stage gyrification proceeds until well after birth. Therefore, there is a clear neurophysiological motivation to understand the physics governing cortical folding and the conditions for normal or pathological cortical folding. Incompatible morphoelastic growth in the cortical layer results in circumferential compression and causes an elastic buckling bifurcation. It is then followed by extreme strains leading to highly folded structures in the post-bifurcation regime. While a theory of axonal tension had been advanced to explain cortical folding under forces imposed by interconnected neurons [12], subsequent studies of cutting followed by elastic relaxation on ferret brains established that axonal tension does not cause folding, while computational studies strongly suggested that incompatible growth does [44]. Bayly et al. [3] explained gyrification patterns by analytic and computational studies based on incompatible morphoelastic growth and Tallinen et al. [38] used experiments in a surrogate, polymeric gel model combined with nonlinear finite element computations to further support the morphoelastic theory of growth .^1^

Wrinkling of surfaces, such as seen on the cortex, and of interfaces, is a common phenomenon. In some cases it is influenced by mismatched elastic moduli between a thin elastic layer and an underlying substrate, a setting common to biological and non-biological thin films [46]. Among the former, it also may control the patterns of wrinkling of fruit and vegetable skins [45]. However, the essence of the phenomenon of brain folding does not depend on stiffness contrasts [32, 10, 11]; the Young’s Modulus of cortical grey matter and of the white matter underlying it are of the same order of magnitude [7, 43]. Therefore, the elastic matter of the folding brain may be reasonably taken as homogeneous.

A number of recent studies have sought to explain aspects of brain folding by incompatible growth under linearized and, more appropriately, nonlinear morphoelasticity [3, 38, 4, 6, 7, 9, 20, 23, 37, 39]. While drawing upon insight from linearized buckling of beams, plates and shells [20, 23, 8], most of the computational work is based on finite strains in the post-bifurcation regime on analytic ellipsoidal shapes. This body of work has shed light on the mechanical conditions governing the development of the organ-wide pathologies of polymicrogyria, pachygyria and lissencephaly [6, 20, 9].

It is notable that the early-forming primary sulci and gyri in humans and other gyrencephalic species show a remarkable robustness of placement in normally developed brains [41]. This is emphasized in fetal brain atlases with data on the geometry of developing brains, such as those obtained from 67 individuals by Gholipour *et al*. [18]. After uniform scaling to normalize volumes, an “average” brain defined by computing the mean geometry showed well-resolved primary folds. This suggests that, when scaled for volume, the placement of those folds is consistent across individuals. Absent this persistence, the folds would have been smeared out in the averaged geometry. A second observation is that despite the organ scale lateral symmetry of the brain the sulci and gyri do not localize into symmetric modes of folding at all scales [22, 30] as seen in computational studies on high-symmetry reference shapes. These observations serve as motivations to identify the sequence of kinematic and mechanical steps that lead to precise placement of the primary folds as well as the range of variation in secondary and tertiary folds. Recent work studied the mechanisms of cell growth and migration and linked them to the developing pattern of the early folds [39, 35]. Here, we note that migration, which is largely complete by around week 26 [41], is responsible for the placement of cells in six layers of the cortex, with later generations occupying outer positions. In turn, this positioning has an influence on subsequent growth and folding.

Here, we take a broader view, seeking to deduce the local volume changes that develop throughout the brain and drive its expansion as well as folding by incompatible, morphoelastic growth. Our approach is a data-driven one. Using magnetic resonance imaging (MRI) data on the geometric changes of the fetal brain, recorded weekly, we seek to solve a series of inverse problems to arrive at the spatially varying growth deformation gradient tensor of the morphoelastic theory. The methods we use begin with MRI segmentation and computational mesh generation to enable image registration across successive weeks of brain development. These steps, themselves involving inverse modelling, provide us with the geometric data for the final stage of physics-constrained inference. Here, we combine direct and adjoint methods for computing gradients of objective functions in a generalized optimization setting, subject to the constraint imposed by the physics of morphoelastic growth. This will leave us with mechanics-constrained geometric data in the form of the precisely defined growth deformation tensor that describes the three-dimensional development of the fetal brain. From this basis, further physically well-founded inference will be possible on the dynamics of fetal brain development. In related work, Garcia et al. [13] used data from fetal MRI studies over weeks 27-37 of development, and anatomical multimodal surface matching to deduce spatiotemporal variations in surface growth. The minimization of the elastic strain energy is used in their image registration approach.

The morphoelastic growth model is discussed in *§*2, the inverse problem for the growth deformation tensor and tests with synthetic data appear in *§*3. MRI segmentation of fetal brain atlas data and computational mesh generation with it appear in *§*4. The MRI registration problem is discussed in *§*5, and the extraction of morphoelastic growth deformation data in *§*6. Results for the inferred growth deformation gradient tensor are in *§*7, and conclusions in *§*8

## 2 The theory of morphoelastic growth

The theory of morphoelastic growth is well-established and traces its roots to multiplicative plasticity, and even before that to multiplicative theormoelasticity. For a discussion of the kinematics we direct the reader to Ref. [14], to Refs [15, 28] for its coupling with mass transport, and to Ref. [2] for a perspective of growth and remodelling. A complete treatment that includes the mathematical background and a proper placement of the theory within nonlinear elasticity can be found in Ref. [19]. The treatment that follows here is rigorous, but eschews formalism in favor of accessibility of the important ideas.

Given the displacement field ***u*** *∈* ℝ^3^, and the reference position of material points ***X*** *∈* ℝ^3^, the deformation gradient tensor is ***F*** = **1** + *∂****u****/∂****X***, where **1** is the isotropic tensor. The multiplicative decomposition of ***F*** that underlies the theory splits it into elastic and growth components, ***F***^e^ and ***F***^g^, respectively, so that ***F*** = ***F***^e^***F***^g^. Incompatibility is admitted by this decomposition in that ***F***^g^, which we think of as driving morphoelastic growth, is not, in general, obtained as a gradient field in the manner that ***F*** arises from ***u***. It therefore does not satisfy the classical kinematic compatibility conditions that ***F*** does.

As explained in the Introduction, we work within the theory of hyperelasticity. We adopt a neo-Hookean strain energy density function *ψ* [39], which depends exclusively on the elastic right Cauchy-Green tensor 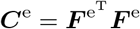,

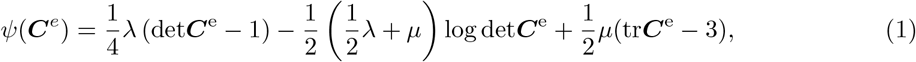

where *µ* and *λ* are the standard Lamé parameters. The first Piola-Kirchhoff stress tensor ***P*** follows as the derivative of the strain energy *ψ*:

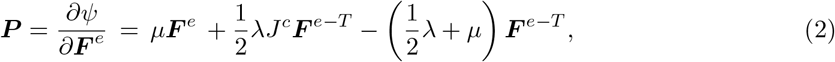

where *J*^c^ = det***F***^*e*^. The first Piola-Kirchhoff stress is governed by the quasistatic balance of linear momentum with no body force:

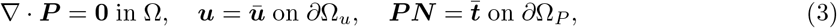

where Ω ⊂ R^3^ denotes the domain, which is the brain, and its Dirichlet and Neumann boundaries are Ω_*u*_ and Ω_*P*_, satisfying 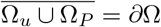 and Ω_*u*_ ∩ Ω_*P*_ = ∅. In this study, 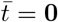 and deformation will be driven by ***F***^g^. As alluded to above, this theory will be cast in the framework of an inverse problem by seeking to match the resulting displacement field with observed data.

### 2.1 A theory of evolving reference configurations

The theory of morphoelastic growth has traditionally been applied to a fixed reference configuration, relative to which the tensors ***F*** and ***F***^g^ have been defined. For finite growth and morphogenesis, which characterise fetal brain development, however, this theory proves inadequate. Its premise is that the entire path of growth and morphogenesis can be described kinematically with the initial state of the brain as the reference configuration. This assumption proves problematic when taken to the logical conclusion that the reference configuration is therefore the singularity when the first brain cell appears. The total growth at all times *t* is ***F***^g^(*t*) relative to this fixed reference configuration with initial condition ***F***^g^(0) = **1**. This hypothesis leads to unphysically large elastic and growth distortions for later times *t* » 0. In numerical implementation of the theory, solvers fail to converge for these large distortions. Furthermore, this theory does not account for mass appearing at some time, say *τ >* 0, thereby introducing material points where none existed before and defining the reference state from which the newly formed material deforms. Finally, it does not address the evolution of local material properties, in this case represented by the strain energy density function. In the traditional approach to morphoelastic growth the strain energy density is defined with respect to the reference configuration. If the latter is fixed, it restricts the changes of the strain energy density function as growth and morphogenesis proceed.

To circumvent these difficulties, we define a continuously evolving reference configuration, Ω_*τ*_, which coincides with the deformed configuration resulting from all morphoelastic processes from times *t* ≤ *τ* (see Figure 1).

**Figure 1.**
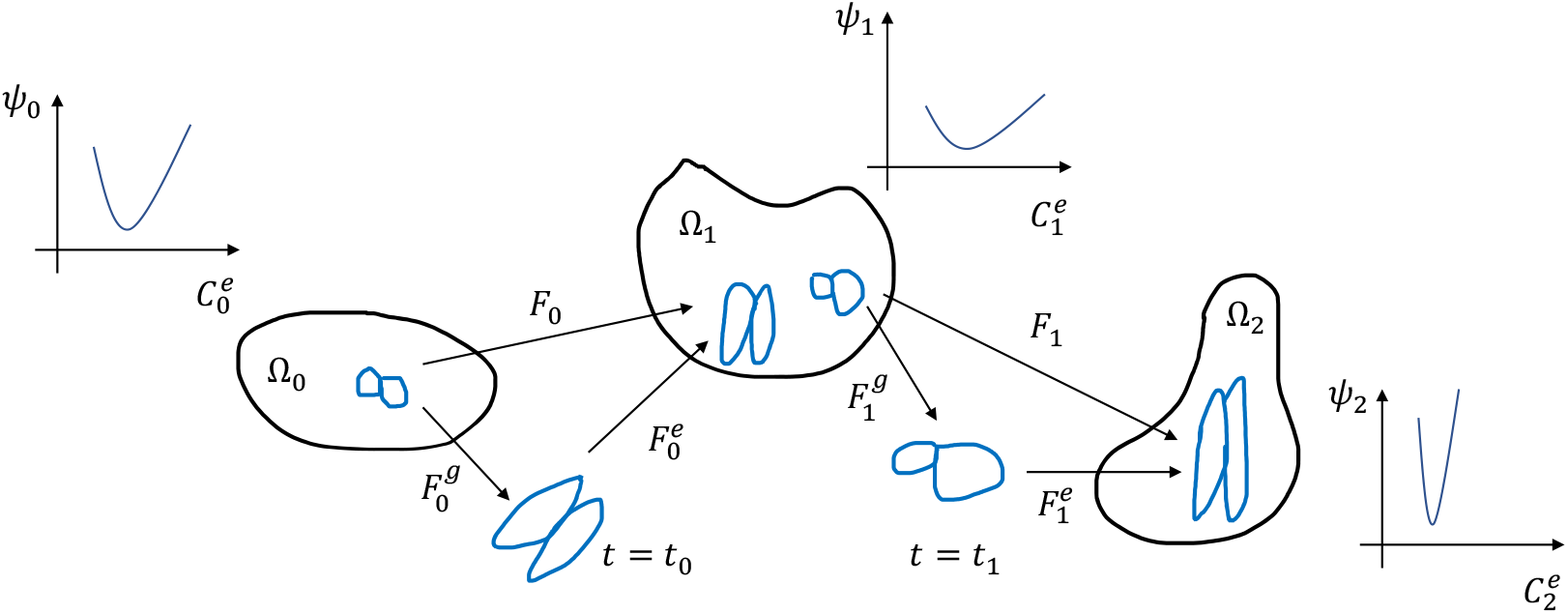
Classical morphoelastic growth presumes that the entire path of growth and morphogenesis can be described kinematically with the initial state of the brain as the reference configuration. In the case of fetal brain development and the emergence of new material, we posit that this assumption proves problematic and propose a theory of evolving reference configurations. Specifically, we split the growth path into multiple individual steps defined by their own reference configuration Ω_*τ*_, kinematics and strain energy density functions defined on them.

In this setting, the kinematics of finite strain multiplicative morphoelasticity is elaborated upon by time parameterization yielding 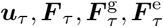 all corresponding to the reference configuration Ω_*τ*_.

They satisfy the kinematic relations:

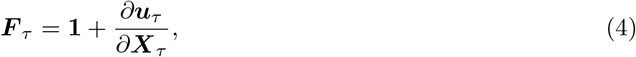

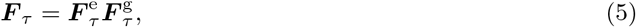

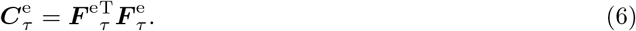

The strain energy density function is defined at points ***X***_*τ*_ *∈* Ω_*τ*_ and written as

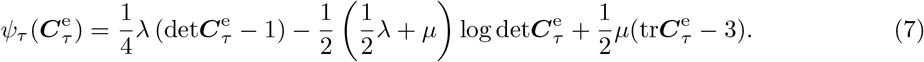

Finally, the stress and governing partial differential equation are:

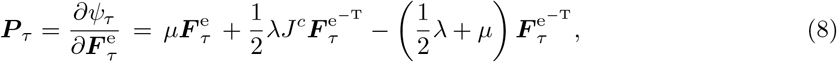

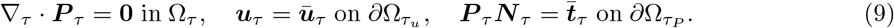

The strain energy density, *ψ*_*τ*_, while written here in time-independent functional form, could also evolve in general. This reflects the understanding that the strain energy density, like the free energy, is defined relative to some reference. Here, as Figure 1 suggests, it is redefined at each reference state, Ω_*τ*_. In practice, a discrete time parameterization is adopted at instants *τ ∈*{*t*_0_, *t*_1_,*…*}. This is natural for data acquisition and computations.

## 3 An inverse problem posed on the geometry of the developing brain

In *§* 4–6 we describe the steps by which we arrive at geometric field data, ***û*** that represents displacements during growth of the developing brain. With these data, we seek to solve an inverse problem for the growth tensor field 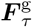 and displacement field ***u***_*τ*_ such that the error ***û***_*τ*_ *–* ***u***_*τ*_ is minimized under the constraint of the physics expressed in Equations (4–9). The data field ***û***_*τ*_ will be interpolated from pointwise displacement vectors 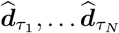 at *N* instants ***X***_*τ*_1, … ***X***_*τ*_*N*. Similarly, we will use the finite-dimensional version 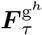 of the unknown growth tensor and the corresponding displacement field, 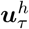. Our approach is to use the finite-dimensional weak form of the governing equations (9), which is expressed as follows in terms of 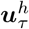 and 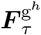:

For some 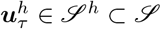, where 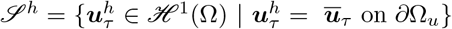, and ∀ ***w***^*h*^ *∈ 𝒱*^*h*^ ⊂ *𝒱*, where *𝒱*^*h*^ = {***w***^*h*^ *∈ℋ*^1^(Ω) | ***w***^*h*^ = 0 on *∂*Ω_*u*_}, the finite-dimensional (Galerkin) weak form of the problem is satisfied:

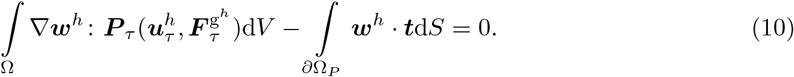

In a forward solution of the weak form, a constitutive model would be written for 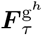. This approach, with some variations, has been followed almost universally in the literature up to this point [3, 38, 4, 6, 7, 9, 20, 23, 37, 39]. The determination of 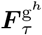 by solution of an inverse problem is a significant departure in the current work. By seeking to invert a tensor field it also stands in contrast to classical inverse problems in mathematical physics that infer a small number of scalar parameters. We decompose Ω_*τ*_ into element sub-domains 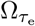, for *e* = 1, … *n*_el_. The variations ***w***^*h*^, trial displacement solutions ***u***^*h*^ and growth tensor 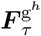 are defined by using a finite number of basis functions in each element,

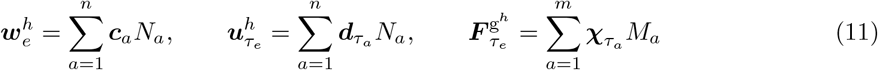

where 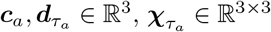, *n* is the dimensionality of the function spaces *𝒥*^*h*^ and *𝒱*^*h*^, *m* is the dimensionality of the expansion for 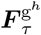 and *N* _*a*_, *M* _*a*_ represent basis functions. We assume the growth tensor to be diagonal and anisotropic, and interpolate it using nodal basis functions, thus reducing its dimensionality also to *n*. Its diagonal terms are written as:

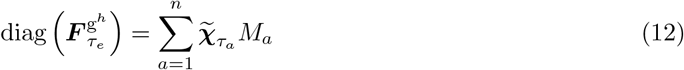

where 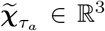. This form was motivated by the total deformation gradient tensor, which when extracted from MRI data on normative, developing fetal brains in Gholipour *et al*.’s atlas [18] by the methods in *§*4–6, was found to be similarly diagonally dominant and anisotropic. We made this assumption throughout the following of this communication, and dispensed with the tildes on ***χ***_*a*_.

We define the residual vector arising from finite element assembly of the weak form:

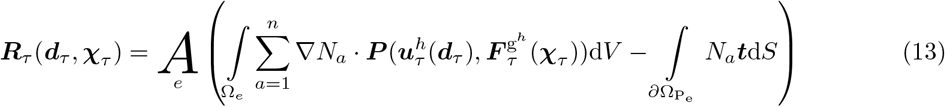

where *A*_*e*_ denotes the assembly operator over the elements, and the arbitrariness of the degrees of freedom corresponding to the variations has been used, as is the practice in the variational setting. Recall that the dimensionality of the vector ***R***_*τ*_ is the total number of unknown displacement degrees of freedom. The discretized, Galerkin weak form of the problem is then ***R***_*τ*_(***d***_*τ*_, ***χ***_*τ*_) = **0**. In the current setting, it represents the physics that constrains the inverse problem, for whose solution we adopt two approaches.

### 3.1 Inverse solution for 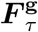 by gradient descent on a loss function

In this approach we directly define the field data ***û*** as a finite-dimensional function:

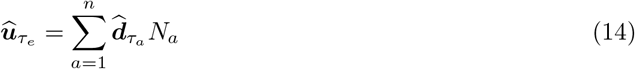

and use it instead of 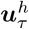 in the weak form (10) and residual equation (13) to arrive at 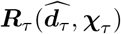. The loss function is

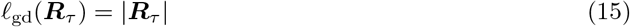

defined via the Euclidean norm. We use gradient descent algorithms, and their variants, to find

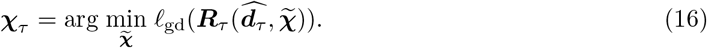

Note that the form of the loss, *𝓁*_gd_ = |***R***_*τ*_ |, means that the exact satisfaction of the constraint 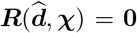 is the optimal solution to (16). As in many high-dimensional, nonlinear optimization problems, this solution is not attainable, in general. Instead, we seek to arrive at *𝓁*_gd_ < *ε* for some tolerance *ε* using either the classical gradient descent algorithm or one of its variants. The field 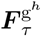 is then recovered by Equation (12).

### 3.2 Solution of the inverse problem by adjoint-based gradient optimization

With ***û***_*τ*_ written as in Equation (14) we solve the following minimization problem, beginning with the loss redefined as the *L*^2^-norm of the error

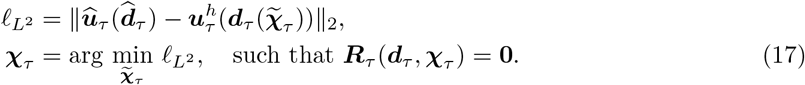

The minimization is solved classically, by computing gradients of the loss 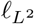. Importantly, the PDE constraint ***R***_*τ*_ (***d***_*τ*_, ***χ***_*τ*_) = **0** makes ***d***_*τ*_ an implicit function of ***χ***. This makes the functional derivatives 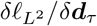 challenging to compute. The obvious approach is to solve the PDE constraint repeatedly for a range of values of ***χ***_*τ*_ and construct the implicit derivative by numerical differentiation. In addition to the expense of a large number of PDE forward solves for a single derivative evaluation, numerical differentiation is noisy and ultimately introduces instabilities. The well-established alternative is to employ the adjoint of the Jacobian of the PDE constraint with respect to ***d***_*τ*_ to compute 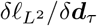. The Jacobian arises in the complete first-order Taylor expansion of the PDE constraint equation, and allows the computation of 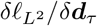 with a single adjoint solution per step. We have adopted this approach to PDE constrained optimization here, and refer to it as *adjoint-based gradient optimization*. In this work we use the *L-BFGS-B* optimization algorithm from *SciPy* [40] with the aid of the *dolfin-adjoint* [27] package for adjoint-based gradient optimization.

### 3.3 Algorithm testing with synthetic data

The gradient descent and adjoint-based gradient optimization approaches were first tested against synthetic data for nonuniform but continuous growth tensor fields. These fields were obtained by solving a three-dimensional, steady state diffusion problem for a scalar field *c* and defining ***F***^g^ to be a function of this argument. The steady state diffusion problem is:

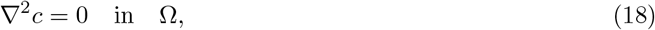

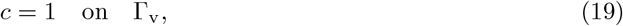

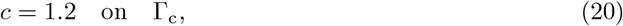

where Ω was taken as the normative fetal brain geometry at week 21 from the atlas of Gholipour *et al*. [18], Γ_v_ is the interface between the ventricles and sub-cortex and Γ_c_ is the outer surface of the cortex. The growth deformation gradient tensor is chosen to be diagonal, but anisotropic, and of the form:

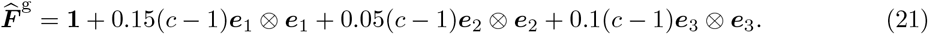

The field of 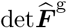 is shown in Figure 2. The forward problem of morphoelastic growth, described in Equations (4–13) was then solved by the finite element method for time *τ* = 0 on a mesh with 27306 tetrahedral elements using the `FEniCS` open source code [1]. The neo-Hookean strain energy density function (1) was used in the the nearly incompressible limit with *λ* = 82200 Pa and *µ* = 1677 Pa [39], corresponding to a Poisson ratio *ν* = 0.49 in the infinitesimal strain regime. We denote the resulting synthetic displacement field by ***u***_s_. To model the noise present in the displacement fields extracted from the fetal brain atlas, varying amounts of Gaussian noise were applied to the synthetic data. For displacement fields with applied noise fraction *p*, nodal displacements were offset by *δ****u*** ∼ *N* (0, *p****u***_*c*_), where *p* ∈{0, 0.01, 0.02}. We did not include spatiotemporal variations of material properties. The results presented here do not change significantly for variations of the bulk and shear modulus within an order of magnitude. However, as is well understood, forward and inverse solution steps converge more slowly as the incompressible limit is approached (studies not shown).

**Figure 2.**
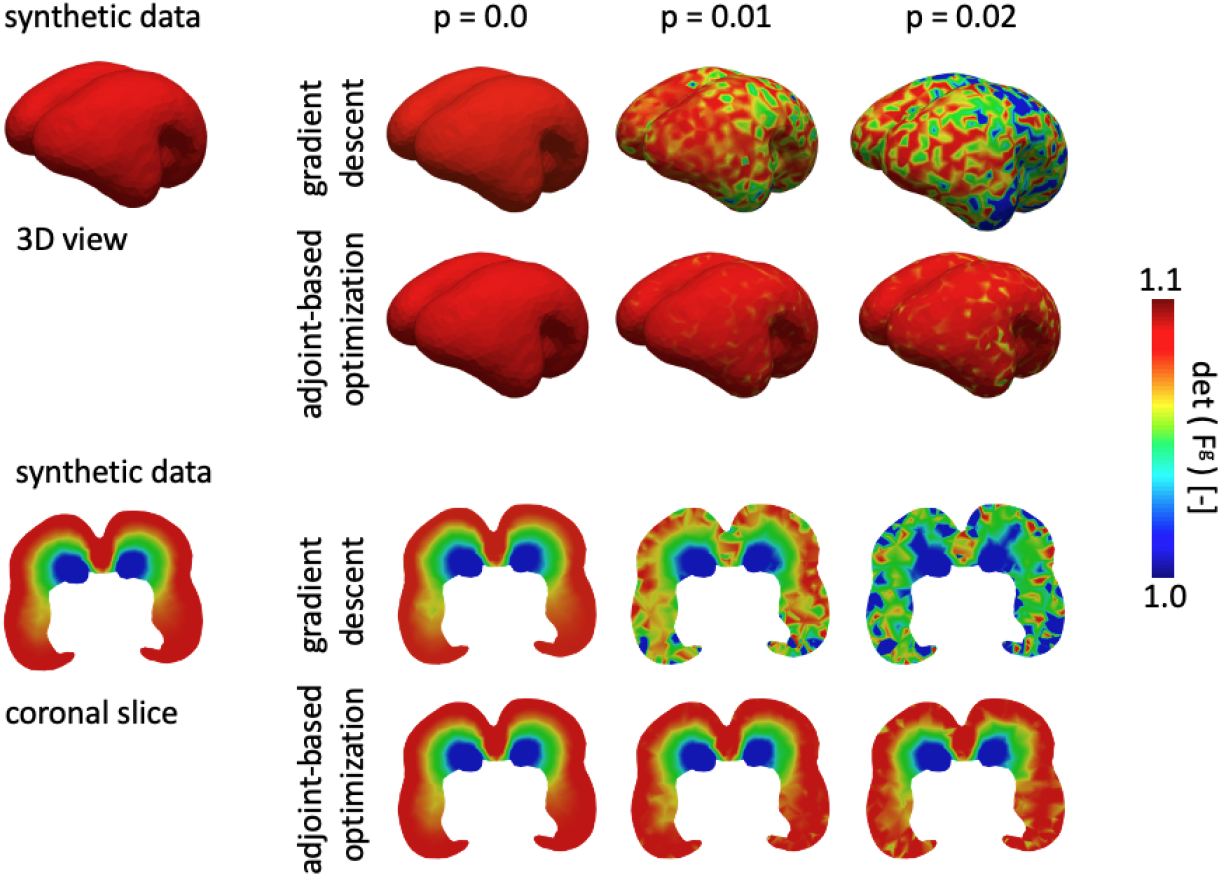
We generated synthetic displacement fields on the fetal brain mesh at 24 weeks in order to test the accuracy of our two optimization algorithms. We show the solution of the inverse problem in the form of the inferred det ***F***^*g*^ fields using the gradient descent and the adjoint-based optimization approach. Top rows show the three dimensional view and the bottom rows shows the coronal view for three levels of superposed noise *p*.

#### 3.3.1 Inverse solution by gradient descent on synthetic data

The problem as posed in *§*3.1 and 3.2 admits a multitude of feasible solutions, and optimal solutions, if they are obtained also could be non-unique. This situation is typical of inverse problems. The optimization algorithms navigate a high-dimensional landscape of feasible solutions seeking the optimal one. Furthermore, stiffness is induced by the nonlinearity of the PDE constraint in the form of the residual (13). This combination can lead to slow convergence or even divergence. Aiming to mollify this problem, we linearly subdivide the synthetic data ***u***_s_ into some number of steps, in this case ten. The gradient descent approach at step *i* uses 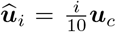. The initial guess for the nodal values of ***F***^g^ at step *i* ≠ 0, i.e., the nodal tensor unknowns ***χ***_*i*_, was the inferred ***F***^*g*^ from the previous step such that 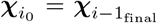. The initial guess for ***χ*** at Step 1 was chosen to be the diagonalized deformation gradient tensor constructed from the displacement 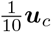, i.e. the target displacement at Step 1. Specifically, we project diagonal components of the deformation gradient tensor to the nodes by solving an *L*^2^ projection problem:

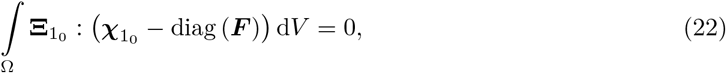

with 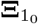 being the variations on 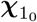. We define the volume averaged *L*^2^ error for the final inferred ***F***^*g*^ at the tenth step as:

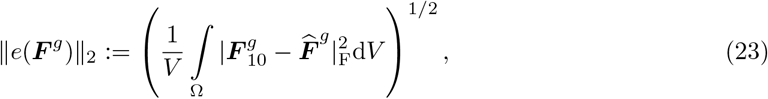

where | · |_F_ denotes the Frobenius norm and 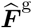 is the field from Equation (21). With the inferred 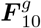, we then evaluate the displacement by solving the forward elasticity problem, and evaluate its volume averaged *L*^2^-error by

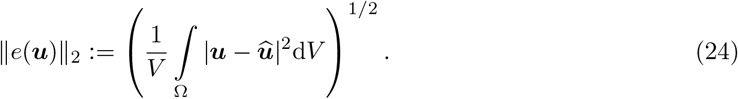

Gradient descent updates were driven by the Adam optimizer with default parameters [26]. For steps 1-9, 10,000 epochs were used with learning rate decay in epoch *k* given by 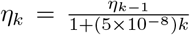 and *η*_0_ = 10^*–*4^. To ensure convergence on the final step, 100,000 epochs were used with learning rate decay given by 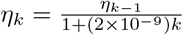 and *η*_0_ = 10^*–*3^.

Figure 2 shows det 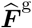 inferred by gradient descent from synthetic data at different level of noise. Also shown in Table 1, gradient descent with the Adam optimizer allows inference of an ***F***^g^ field with volume-averaged *L*^2^-error that is three orders of magnitude lower (fifth column) than the *L*^*∞*^-norm of the applied ***F***^g^ for synthetic data generation, and even with noise fraction *p* = 0.02 remains an order of magnitude lower. However, the inferred ***F***^g^ field appears less smooth when obtained from the noisy data. We also have included the volume-averaged *L*^2^-error in the forward displacement (fourth column) computed by applying the gradient descent-inferred ***F***^g^ field. For synthetic data without noise, the volume-averaged *L*^2^-error is three orders of magnitude lower than the *L*^*∞*^ -norm of ***û*** and one order of magnitude lower for noise with *p* = 0.02.

**Table 1:** Results of the inverse problem for synthetic data. The superscripts “gd” and “adj” denote the solution obtained via using gradient descent with the Adam optimizer and adjoint-based optimization, respectively.

| Noise, $p$ | $\ \hat{\mathbf{u}}\ _\infty$ | $\ \mathbf{F}^g\ _\infty$ | $\ e(\mathbf{u})\ _2^{\text{gd}}$ | $\ e(\mathbf{F}^g)\ _2^{\text{gd}}$ | $\ e(\mathbf{u})\ _2^{\text{adj}}$ | $\ e(\mathbf{F}^g)\ _2^{\text{adj}}$ |
| --- | --- | --- | --- | --- | --- | --- |
| 0.00 | $5.212 \times 10^{-1}$ | 1.767 | $6.697 \times 10^{-4}$ | $1.640 \times 10^{-3}$ | $6.415 \times 10^{-5}$ | $5.148 \times 10^{-3}$ |
| 0.01 | $5.196 \times 10^{-1}$ | 1.767 | $9.978 \times 10^{-3}$ | $7.352 \times 10^{-2}$ | $5.476 \times 10^{-4}$ | $1.053 \times 10^{-2}$ |
| 0.02 | $5.284 \times 10^{-1}$ | 1.767 | $1.753 \times 10^{-2}$ | $1.183 \times 10^{-1}$ | $1.104 \times 10^{-3}$ | $2.001 \times 10^{-2}$ |

#### 3.3.2 Inverse solution by adjoint-based gradient optimization

As discussed in *§*3.2 adjoint-based gradient optimization involves the forward solution of the PDE constraint ***R***(***d, χ***) = **0** at each step of the algorithm–see Equation (17). As also expressed there, this forward solution is driven by the inferred nodal growth tensor field ***χ*** at each iteration. This forward problem is numerically stiff due to the nonlinearity, near-incompressibility and complex geometry of the brain. While the adjoint solution step to determine gradients typically poses no difficulty, divergence of the forward solution will cause the termination of the overall algorithm. Therefore, we now linearly subdivide the inferred ***χ*** into 100 steps in driving the forward solution. The initial guess for ***χ*** was again chosen to be the diagonalized deformation gradient tensor constructed from ***u***_s_.

Figure 2 and Table 1 also include the results obtained by adjoint-based gradient optimization. Using noise-free data, the volume-averaged *L*^2^-error in the inferred ***F***^*g*^ is higher than that obtained by the gradient descent approach, but the volume-averaged *L*^2^-error in the forward displacement solution obtained as an inherent part of the adjoint-based gradient optimization approach is about one order of magnitude lower than the corresponding error obtained by gradient descent. The superiority of the adjoint-based approach is more apparent in the presence of noise, improving to an order of magnitude lower volume-averaged *L*^2^-error for ***u*** and ***F***^g^ over the gradient descent approach for *p* = 0.02. Additionally, the inferred ***F***^g^ field is smoother than that obtained by gradient descent. The adjoint-based approach is, in general, more computationally expensive since it needs one evaluation of the adjoint solution per step. Nevertheless, given these performance metrics, we choose to exclusively use the adjoint-based gradient optimization approach with the real MRI data from the fetal brain atlas, because of the inevitability of noise therein.

We emphasize that the above numerical implementations of the inverse problem was made possible by the theory of evolving reference configurations. The traditional approach of a single reference configuration with ***F*** (*t*) = ***F***^e^(*t*)***F***^g^(*t*) does not converge for times beyond week 23.

## 4 MRI Segmentation and FE Model Generation

We obtained data on brain geometries from a spatiotemporal magnetic resonance imaging (MRI) atlas of the fetal brain developed for the study of early brain growth [17]. Based on MRIs of 81 normal fetuses scanned between gestational weeks 19 and 39, Gholipour *et al*. created a four-dimensional atlas of brain development during the second half of gestation and covering weeks 21 through 37 [18]. Six to eight scans were used for the reconstruction of each week’s atlas. The automatic atlas generation includes repeated motion correction, super-resolution volume reconstruction, brain mask segmentation, rigid alignment to the atlas space and intensity homogenization [17]. The resulting atlas clearly illustrates the temporally and spatially heterogeneous growth during early *in utero* brain development, including numerous instances of folding and creasing. In a first step, the present work focuses on weeks 21 through 25 during which the first major elastic bifurcation occurs, and from which the central sulcus (CS) emerges [21]. For each gestational week, we created a finite element model of the brain from the respective MR images using the `ScanIP` software environment of Simpleware (Synopsys, Mountain View CA), see Figure 3. In a semi-automatic segmentation procedure, we delineated the cortex, subcortex and lateral ventricles based on grayscale contrast and created a three-dimensional reconstruction of these structures [42]. The software converted these segmentations into a volumetric model consisting of tetrahedral elements. We prescribed a minimum and maximum element edge length of 2.0mm and 2.5mm, respectively, and obtained meshes with a total number of 68849 elements for the model of week 21, 78385 elements for week 22, 83000 elements for week 23, 97138 elements for week 24 and 172289 elements for week 25. The number of elements and nodes of each subregion are summarized in Table 2. Based on our segmentations, we observe that the total brain volume, i.e. cortex and subcortex, increases by 130% and ventricular volume increases by 17% between weeks 21 and 25. Specifically, cortical volume changes from 17155 mm^3^ at week 21, to 20651 mm^3^ at week 22, 20468 mm^3^ at week 23, 26099 mm^3^ at week 24 and 35232 mm^3^ at week 25; subcortical volume increases from 23834 mm^3^ at week 21, to 29159 mm^3^ at week 22, 33056 mm^3^ at week 23, 41360 mm^3^ at week 24 and 58646 mm^3^ at week 25; and ventricular volume changes from 5079 mm^3^ at week 21, to 5176 mm^3^ at week 22, 4011 mm^3^ at week 23, 4527 mm^3^ at week 24 and 5946 mm^3^ at week 25. The rostral-caudal brain length increases by 29.5% between weeks 21 (59.86 mm) and 25 (77.54 mm); the width of the brain increases by 26.4% between weeks 21 (49.31 mm) and 25 (62.33 mm); and the height of the brain increases by 33.8% between weeks 21 (39.19 mm) and 25 (52.44 mm).

**Figure 3.**
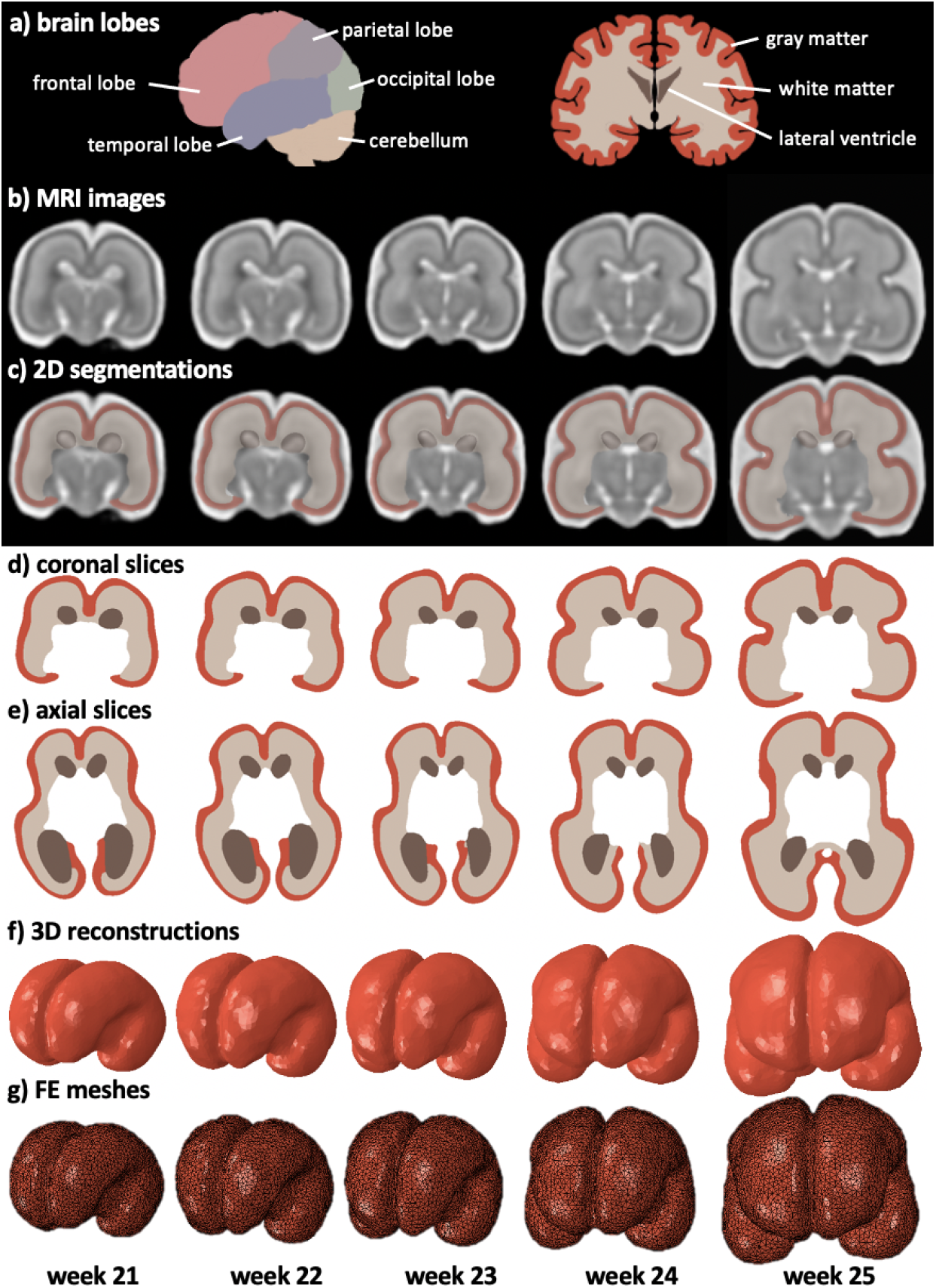
Brain anatomy, MR image segmentation, and finite element model generation. (a) The brain consists of frontal lobe, temporal lobe, parietal lobe, occipital lobe, and the cerebellum. The cerebrum can be separated into cortical gray matter and subcortical white matter layers, as well as the fluid filled lateral ventricles. We use (b) structural MRIs from gestational weeks 21 through 25 and create 3D reconstructions based on a semi-automatic segmentation process (c). In (b) We delineate the cortex, subcortex and ventricles based on their grayscale thresholding and manual correction. (d-e) show coronal and axial slices of the segmentation, respectively. The fully (f) three-dimensional reconstructions are converted into (g) volumetric finite element models that consist of tetrahedral elements.

**Table 2:** Mesh properties of our five finite element models for gestational weeks 21 through 25. Proportional to the increase in mesh size, total brain volume increases by 130% during this time period. #e = number of elements; #n = number of nodes.

| model | week 21 |  | week 22 |  | week 23 |  | week 24 |  | week 25 |  |
| --- | --- | --- | --- | --- | --- | --- | --- | --- | --- | --- |
|  | #e | #n | #e | #n | #e | #n | #e | #n | #e | #n |
| cortex | 25747 | 8262 | 29363 | 9439 | 29873 | 9623 | 34618 | 11188 | 97945 | 14957 |
| subcortex | 35749 | 8712 | 41996 | 9995 | 46852 | 10894 | 55066 | 12911 | 65217 | 15576 |
| ventricles | 7353 | 2106 | 7026 | 2068 | 6275 | 1881 | 7454 | 2157 | 9127 | 2638 |
| total | 68849 | 14262 | 78385 | 16204 | 83000 | 17095 | 97138 | 19982 | 172289 | 25197 |

## 5 MRI Registration Framework

The continuous morphological changes of the fetal brain during *in utero* development are inherently contained in the fetal brain atlas described previously. To determine the incremental brain deformations driven by growth between consecutive gestational weeks, we use a previously developed registration method that determines the non-rigid spatial transformation between two MR images by maximizing the congruence of image intensities. Specifically, we built on the work of Pawar et al. [31] who optimized their algorithm for large deformations and topological changes between medical images. The source image *I*_1_(*f* (***x***, *t*)) and the target image *I*_2_(***x***) are both embedded in hierarchical truncated B-spline objects with the spatial transformation function *f* (***x***, *t*) given by [31]

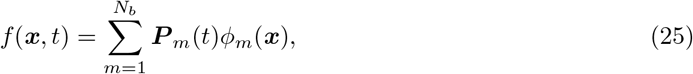

where **P**_*m*_(*t*) are the control points associated with the trivariate basis functions *φ*_*m*_(***x***), and *N*_*b*_ represents the total number of basis functions. As part of the registration process, the transformation function *f* (***x***, *t*) is incrementally varied until dissimilarities between source and target image are minimal. To that end, we followed the proposed energy functional *E*(*f* (***x***, *t*)) proposed by Pawar et al. which accounts for intensity differences and penalizes non-smoothness of the deformation field [31]. The minimization of the energy functional is achieved by posing it as an *L*^2^ gradient flow, thus simplifying the optimization problem to a partial differential equation. The energy functional takes the following form [31]

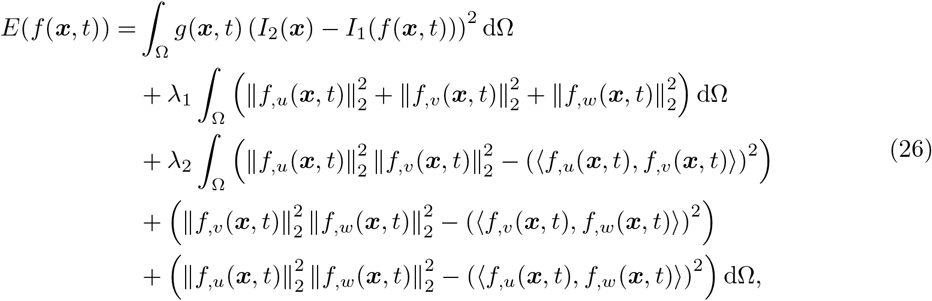

where the first term measures the sum of squared differences of the intensity between the iteratively updated source and target images, ⟨*u, v*⟩ is the dot product and *λ*_1_ and *λ*_2_ are regularization parameters that penalize non-smoothness and inconsistent area change of each face of the 3D control grid elements during deformation. The terms *f*_,*u*_(***x***, *t*), *f*_,*v*_(***x***, *t*) and *f*_,*w*_(***x***, *t*) are the first derivatives of *f* (***x***, *t*) with respect to coordinates *{u, v, w}*, and *g*(***x***, *t*) is given by

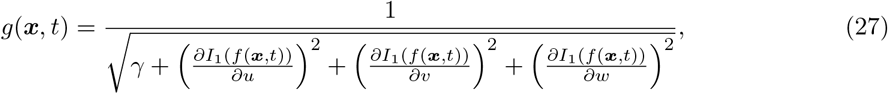

where *γ* is a small number to prevent division by zero. The gradient flow form for minimization of *E* is:

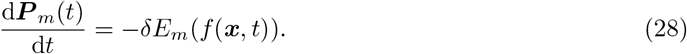

Control points are updated using the Forward Euler method and by introducing a pseudo timestep *ϵ*. The control points *P*_*m*_(*t*) are iteratively computed for time point *s* + 1 based on the solution of the previous timestep *s* as follows

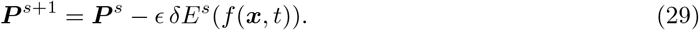

*δE*^*s*^(*f* (***x***, *t*)) is the derivative of the energy functional with respect to parametric domain ***x***, see Ref. [31] for a detailed derivation. The optimization loop ends when the change in intensity difference falls below a given tolerance. We direct the reader to Ref. [31] for a detailed derivation of the registration framework. Also of interest is Ref. [13] where the authors used minimization of the elastic strain energy in their image registration of morphogenetic changes in fetal brains. In the work presented here, we embed our images in an initial three-dimensional grid of size 32 × 32 × 32 control points, set maximum number of refinement steps to 3, regularization parameters *λ*_1_ and *λ*_2_ to 0.0001 and 0.0001, respectively, and chose a timestep size of 1 × 10^*–*5^.

We used the registration framework to determine the four deformation fields between weeks 21 and 22, weeks 22 and 23, weeks 23 and 24 and weeks 24 and 25. For each pair, we selected the first week as the source image and the second week as the target image. It took 22 iterations for the first two steps of 21-22 and 22-23 weeks, 24 iterations for 23-24 weeks and 57 iterations for 24-25 weeks to obtain the optimal transformation map with an average similarity ratio of 81.95%. This increase in iterations reflects the evolving morphological complexity of the progressive developmental steps. Local spline refinement increased the number of active degrees of freedom on average by a factor of 5.3; additional convergence properties are summarized in Table 3.

**Table 3:**
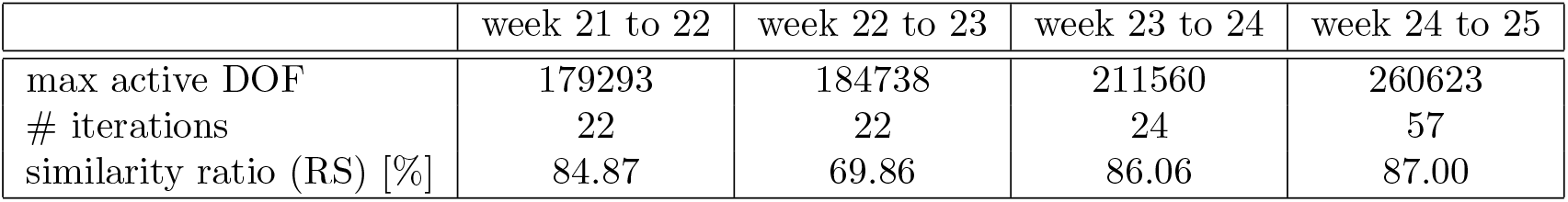
Convergence properties of the registration framework. We achieve a mean similarity ratio of 81.95% after registering two consecutive weeks. Local spline refinement increased the number of active DOFs on average by a factor of 5.3.

|  | week 21 to 22 | week 22 to 23 | week 23 to 24 | week 24 to 25 |
| --- | --- | --- | --- | --- |
| max active DOF | 179293 | 184738 | 211560 | 260623 |
| # iterations | 22 | 22 | 24 | 57 |
| similarity ratio (RS) [%] | 84.87 | 69.86 | 86.06 | 87.00 |

## 6 Growth-Induced Full-Field Brain Deformations

Following the registration step, we extracted the displacement vector of each control point in our grid. In Figure 4 we show the undeformed and deformed grids on a coronal and axial slice for all four registration steps. The effect of the regularization terms is clearly reflected in the smoothness of the deformation field throughout the brain. Simultaneously, the week-wise registration steps allow identification of the major folding event, i.e. the formation of the central sulcus, at week 24. Increased grid density leads to a higher spatial resolution of the three-dimensional deformation field and improves the detection of local growth phenomena.

**Figure 4.**
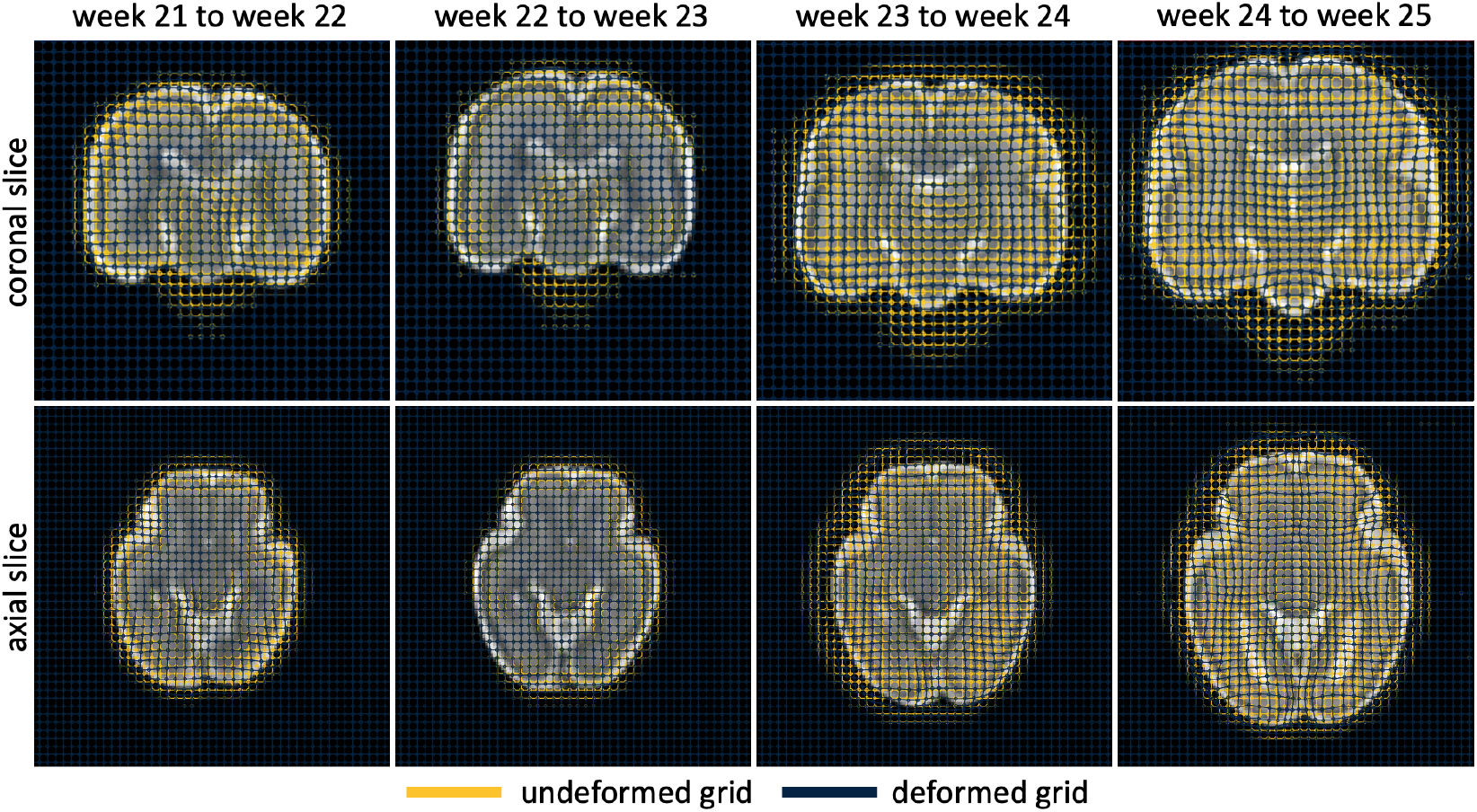
The registration framework iteratively updates the positions of control points that belong to the spline object embedded in each week’s MRI data. Here, we plot the undeformed and deformed grids in representative coronal and axial slices for the registrations between weeks 21 and 22, weeks 22 and 23, weeks 23 and 24 and weeks 24 and 25. From the MRI images we observe the overall volume increase of the brain during the 5 week period. The two grids per image show the increasingly heterogeneous displacement field with rather uniform morphogenetic growth between weeks 21 and 23 and more localized displacement patterns in individual lobes between weeks 23 and 25.

Figure 5 shows registration results for changes between weeks 24 and 25 in three representative slices, the coronal, axial and sagittal views, respectively. The magnified images reveal the grid deformation and identify local growth patterns that produce highly heterogeneous deformation fields. Two challenges are encountered in the steps of MR image registration and computation of growth-induced deformation:

**Figure 5.**
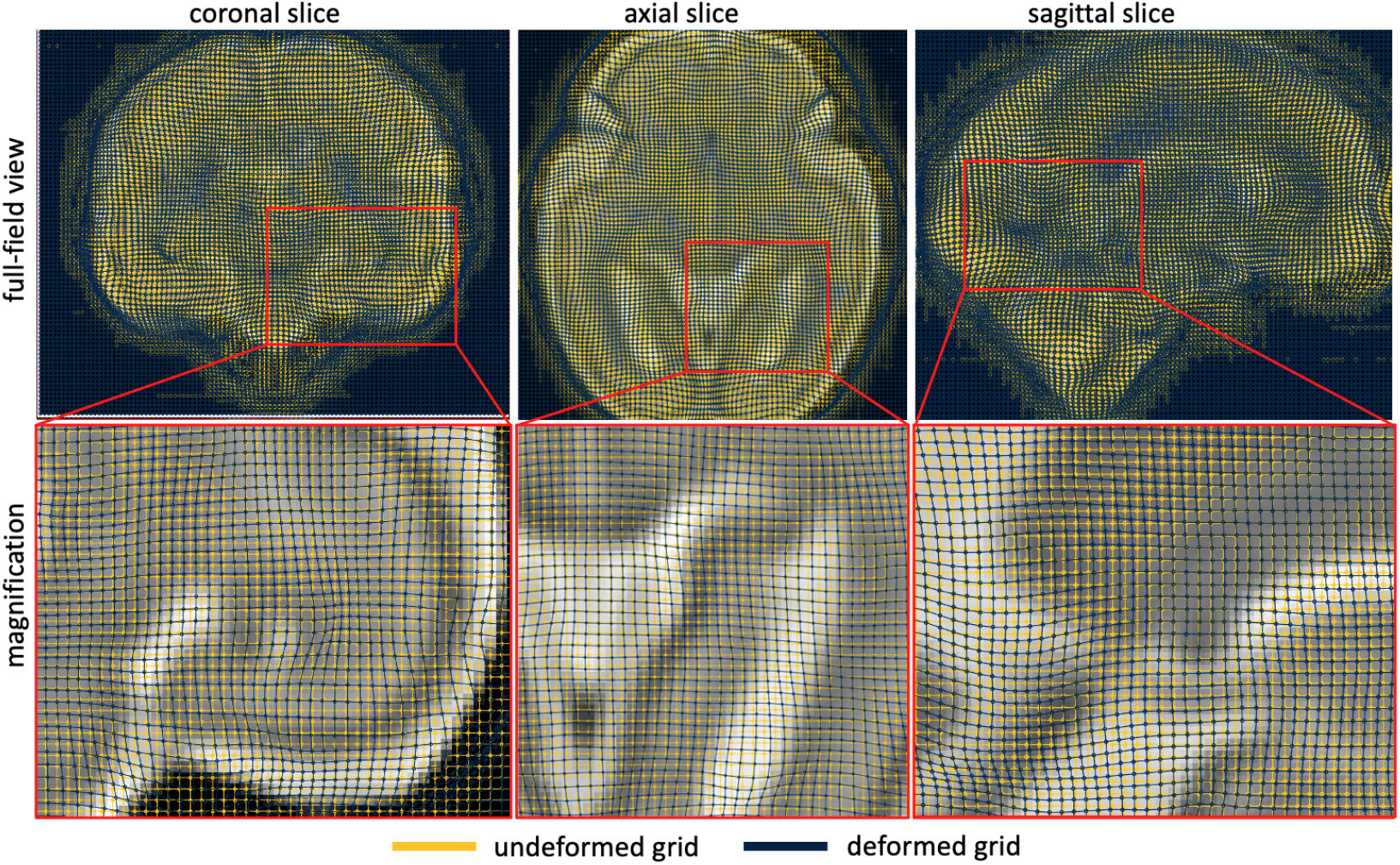
Coronal, axial and sagittal views of the registration results for week 24 to 25. We notice increasingly heterogeneous displacement patterns due to localized growth in distinct subregions of the brain. The registration framework delivers highly smooth displacements of the control points-to the extent that some local phenomena might be eliminated due to over-regularization of the spline object. As a consequence of this smoothing, the emergence of new substructure between two scans will lead to artificial grid distortions. Overall, the registration delivers a reliable displacement field representative of the temporally and spatially varying growth patterns [5].

### Newly formed brain regions limit the registration framework

The registration framework faces significant challenges when new substructures emerge between two distinct scans. In general, the registration framework assumes that all material points are preserved and simply undergo a potentially large deformation. During fetal brain development, novel brain structures emerge between discrete weekly atlases. The generation of new material points leads to non-uniformities and incompatibilities in the displacement field which we have not yet addressed in the present work. Effectively, registration is not being performed on the same brain in two different configurations, but on two brains with different structures. Since our registration framework minimizes pixel intensity differences, the appearance of novel structures, can lead to divergence of our registration step and produce a severely distorted displacement field. Therefore, our approach provides reliable displacement data in the case of morphogenetic growth which manifests in the form of pure volumetric expansion. In this case, material points preserve their intensity value in MR images and simply displace. When new material emerges and influences the intensity distribution, the registration framework is observed to artificially distort the grid. Therefore, we are limited to one-week intervals over which the emergence of new substructures is minimal.

### Heterogeneous Growth Field

In order to prepare data for the inverse problem, we use the nodal displacement vectors in our FE meshes for weeks 21 through 24 to determine the reference configurations. Specifically, we use trilinear interpolation in the registration data to obtain the full-field displacement data for every node in each mesh based on the registration results from that particular week. Figure 6 shows the respective results as displacement vectors that are color-coded by magnitude. Earlier weeks (21 to 23) are characterized by rather homogeneous small displacements across the cortex. Later weeks (23 to 25) exhibit increasingly heterogeneous displacements which is characteristic for localization of growth due to the formation of the central sulcus, and the subsequent formation of folds within each lobe. The rapid proliferation and migration of neurons during this period of development [5] leads to an acceleration of brain growth. Neuronal migration is largely complete by week 26 [41], but their placement has an influence on subsequent growth and folding. The observed growth patterns are also indicative that brain development is significantly more complex than purely uniform, morphological growth but must adhere to genetically encoded cell migration patterns that result in the highly reproducible brain topology observed within any species. The top row of Figure 6 shows the displacement field of the outer cortical surface; the bottom row shows the displacement field of the ventricular surface. We measured a maximum displacement of 5.79 mm in the temporal lobe between weeks 24 and 25. We observe mean displacements of the outer cortical surface of 0.45± 0.27 mm between weeks 21 and 22, 0.88± 0.42 mm between weeks 22 and 23, 1.91± 0.74 mm between weeks 23 and 24 and 3.03± 1.06 mm between weeks 24 and 25. Mean displacements of the ventricular surface are 0.19± 0.13 mm between weeks 21 and 22, 0.28± 0.19 mm between weeks 22 and 23, 0.8± 0.51 mm between weeks 23 and 24 and 1.49± 0.7 mm between weeks 24 and 25. Overall, we find that growth is highly symmetric during this early stage of brain development and posit that individual differences between hemispheres are the result of averaging data from multiple brains when the atlas was constructed [17].

**Figure 6.**
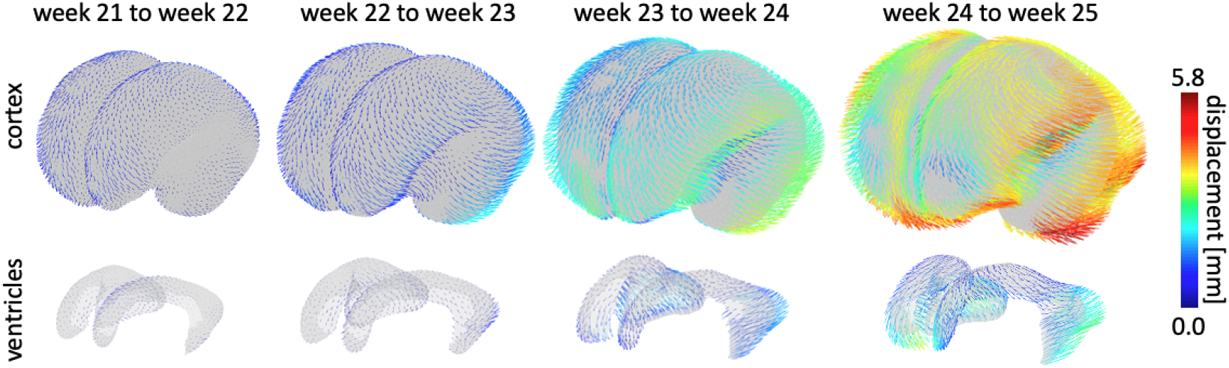
Full-field displacement data between week 21 and 22, week 22 and 23, week 23 and 24 and week 24 and 25. We observe a homogeneous displacement field between weeks 21 and 23 and increasingly heterogeneous displacement patterns between week 23 and 25. Specifically, we notice that the emergence of the central sulcus leads to localization of growth patterns that are attributed to the onset of secondary buckling in individual lobes and increased folding of the cortical surface. A maximum displacement of 1.24 mm was observed from week 21 to 22, 2.29 mm was observed from week 22 to 23, 3.65 mm was observed from week 23 to 24 and 5.79 mm from week 24 to 25.

In earlier work, Rajagopalan and co-workers had performed image registration on fetal brains over weeks 20-28 to report the scalar volume changes [33] as well as the “principal growth direction” [34] and the spatiotemporal variation of these quantities. The same information could have been extracted from the results presented in Figures 4-6. Instead, we seek to further account for the constraint of the physics of morphoelastic growth. As we show in Section 7, this further delineates the inelastic growth deformation tensor, ***F***^g^ and separates it from the elastic part of the deformation gradient tensor, ***F***^e^. This consideration of the laws of morphoelastic growth differentiates the current work from that of Rajagopalan and co-workers [33, 34].

### 6.1 Gaussian Filtering

While filtering techniques typically are applied to raw MRI data to generate the images in Figure 3a, noise is reintroduced by the registration algorithm. The displacement field reported in Figures 5-6 is therefore in need of smoothing before its use in the numerical techniques of inverse modelling, since the forward and inverse problems require computation of deformation gradients. Further Gaussian filtering helps improve convergence of the corresponding numerical solvers (as shown in Tables 4). We applied Gaussian filtering to the post-registration displacement field, noting however, that the standard discrete Gaussian filter cannot be applied in a straightforward manner to unstructured meshes that must be used for the irregular geometry of the brain. Consider the continuous Gaussian filter over the infinite domain:

**Table 4:** Results summary. The values of standard deviation *σ* correspond to Gaussian filtering of the registration data. “N/A” denotes no filters applied. From the triangle inequality, the total volume-averaged *L*^2^-error in the forward displacement field from the inference relative to the MRI displacement data is bounded from above by 4.3 × 10^*–*2^ for either value of *σ*.

| week | $\sigma$ mm | $\ \hat{\mathbf{u}}\ _{\infty}$ mm | $\ e(\mathbf{u})\ _2$ mm |
| --- | --- | --- | --- |
| 21-22 | N/A | $1.820 \times 10^0$ | $1.523 \times 10^{-2}$ |
| | 0.5 | $1.593 \times 10^0$ | $1.177 \times 10^{-2}$ |
| 22-23 | N/A | $1.425 \times 10^0$ | $1.453 \times 10^{-2}$ |
| | 0.5 | $1.307 \times 10^0$ | $1.162 \times 10^{-2}$ |
| 23-23.5 | N/A | $1.451 \times 10^0$ | $2.098 \times 10^{-2}$ |
| | 0.5 | $1.283 \times 10^0$ | $1.773 \times 10^{-2}$ |
| 23.5-24 | N/A | $1.451 \times 10^0$ | $2.154 \times 10^{-2}$ |
| | 0.5 | $1.283 \times 10^0$ | $1.770 \times 10^{-2}$ |
| 24-24.125 | N/A | $6.279 \times 10^{-1}$ | $6.739 \times 10^{-3}$ |
| | 0.5 | $5.454 \times 10^{-1}$ | $5.373 \times 10^{-3}$ |
| 24.125-24.25 | N/A | $6.279 \times 10^{-1}$ | $6.818 \times 10^{-3}$ |
| | 0.5 | $5.454 \times 10^{-1}$ | $5.429 \times 10^{-3}$ |
| 24.25-24.375 | N/A | $6.279 \times 10^{-1}$ | $6.635 \times 10^{-3}$ |
| | 0.5 | $5.454 \times 10^{-1}$ | $5.502 \times 10^{-3}$ |
| 24.375-24.5 | N/A | $6.279 \times 10^{-1}$ | $6.776 \times 10^{-3}$ |
| | 0.5 | $5.454 \times 10^{-1}$ | $5.336 \times 10^{-3}$ |
| 24.5-24.625 | N/A | $6.279 \times 10^{-1}$ | $6.914 \times 10^{-3}$ |
| | 0.5 | $5.454 \times 10^{-1}$ | $5.331 \times 10^{-3}$ |
| 24.625-24.75 | N/A | $6.279 \times 10^{-1}$ | $6.488 \times 10^{-3}$ |
| | 0.5 | $5.454 \times 10^{-1}$ | $5.349 \times 10^{-3}$ |
| 24.75-24.875 | N/A | $6.279 \times 10^{-1}$ | $6.396 \times 10^{-3}$ |
| | 0.5 | $5.454 \times 10^{-1}$ | $5.162 \times 10^{-3}$ |
| 24.875-25 | N/A | $6.279 \times 10^{-1}$ | $6.298 \times 10^{-3}$ |
| | 0.5 | $5.454 \times 10^{-1}$ | $5.078 \times 10^{-3}$ |

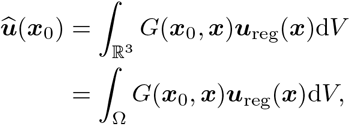

where 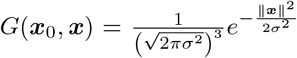 is the three-dimensional Gaussian distribution, *σ* is the standard deviation and ***u***_reg_ is the displacement field after registration. Since ∫_Ω_ *G*d*V <* 1 we scale the filtered displacement at each node to obtain:

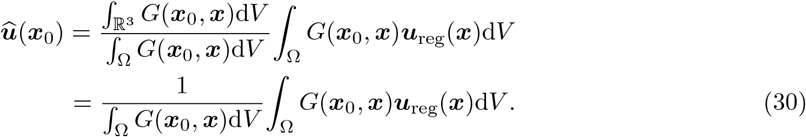

## 7 Inference of the fetal brain’s growth deformation tensor

The displacement field data for weeks *τ* to *τ* + 1 obtained after registration and filtering, as detailed in §4–6, is ***û***_*τ*_. The corresponding nodal values on various meshes are 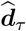. As explained at the end of *§*3.3.2, we used adjoint-based gradient optimization guided by the lower volume-averaged *L*^2^- errors obtained relative to optimization by gradient descent. The following subsections discuss the meshes used, further interpolation of data between 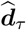 and 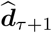 to aid convergence, initialization of ***χ*** (nodal values of ***F***^g^) and numerical performance. Results are presented as tables and figures for the volume-averaged *L* -errors, Equation (24), with ***û*** = ***û***_*τ*_ and figures for the inferred fields of ***F***^g^.

### 7.1 Meshes

The MRI data at weeks 21 and 23 yield the corresponding reference configurations, Ω_21_ and Ω_23_, on which tetrahedral meshes were constructed with 27306 and 32385 elements, respectively. Reference configurations Ω_22_ and Ω_24_ were then generated by deforming Ω_21_ and Ω_23_, respectively, using the displacement fields ***û***_21_ and ***û***_23_ obtained by MRI registration for week 21-22 and week 23-24. These displacement fields applied to the meshes on Ω_21_ and Ω_23_ also yield the meshes on Ω_22_ and Ω_24_. All these meshes appear in Figure 7.

**Figure 7.**
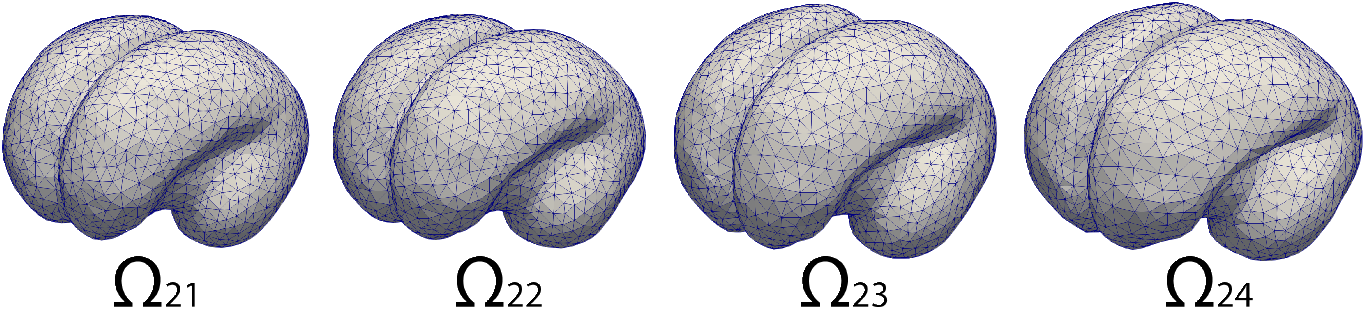
We generated tetrahedral meshes based on the segmentation at each gestational week 21 through 24 in support of our proposed theory of evolving reference configurations. At 24 weeks, the central sulcus begins to emerge and the temporal lobe expands noticeably.

### 7.2 Data interpolation to aid convergence

The displacement field data, ***û***_23_, between weeks 23 and 24, results in large distortions that appear in the deformation gradient, 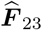. Since this field also drives, and confers these distortions on, the iterates of the inferred 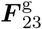, it makes the forward solution for ***u***_23_ numerically stiff and prone to divergence in the adjoint-based gradient optimization. We therefore carried out a linear interpolation and redefined the displacement on Ω_23_ to be 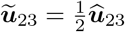 to an interpolated reference configuration Ω_23.5_. In a continuation of this interpolation, we also defined 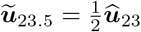 on Ω_23.5_ to Ω_24_. In a further magnification of this large morphoelastic growth, ***û***_24_ between weeks 24 and 25 leads to even greater distortions and more severe divergence of the forward solution for ***u***_24_ during adjoint-based gradient optimization. We therefore defined eight intermediate displacement fields 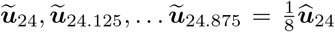 and the corresponding interpolated reference configurations Ω_24.125_, … Ω_24.875_. These interpolated geometries fit with the concept of evolving reference configurations discussed in Section 2.1. Given these interpolated displacement fields, we aimed to infer the growth deformation tensor, 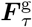, between reference configurations Ω_*τ*_ and Ω_*τ*+∆*τ*_ defined as above.

The initial guess at each configuration 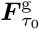 was chosen to be diagonal and assembled from the corresponding components of the deformation gradient tensor 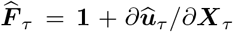 or 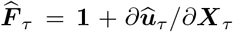(if displacement interpolation to intermediate reference configurations was used). The nodal values, 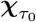 were then obtained by solving the *L*^2^-projection:

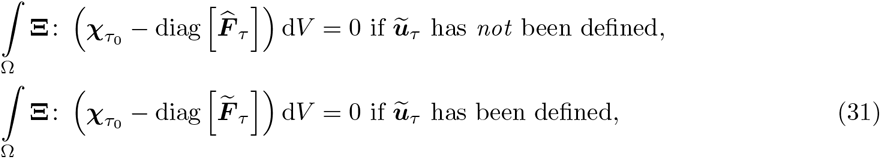

with **Ξ** being the variations on 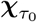.

### 7.3 Convergence

Our approach to the inverse problem involves iterations on the adjoint equation to update ***χ***_*τ*_, which is then interpolated for 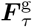. Each solution of the adjoint equation is followed by a forward solution for ***u***_*τ*_. In order to mollify numerical stiffness and ease the direct solver’s path to convergence, we linearly subdivided ***χ***_*τ*_ into 100 steps in driving the forward solution. The convergence threshold was set to requiring that the loss (see Equation 17) be smaller than 2 × 10^*–*2^ of ∥ *û*_*τ*_∥_*∞*_, and that the relative change in loss between successive adjoint solution steps falls below 10^*–*3^. This threshold typically required 2000 adjoint iterations to achieve and cost about 900 CPU-hours on the XSEDE cluster *Comet*. A sample convergence plot is shown in Figure 8. Linearly extrapolating the loss curve, we see that it would take at minimum an additional 1000 iterations to see the loss drop by another order of magnitude, but given the convex shape of this curve, it would likely require many more. Given the computational cost that this would incur, we have not pursued a further decrease of the relative change in loss by an order of magnitude to *∼* 10^*–*4^.

**Figure 8.**
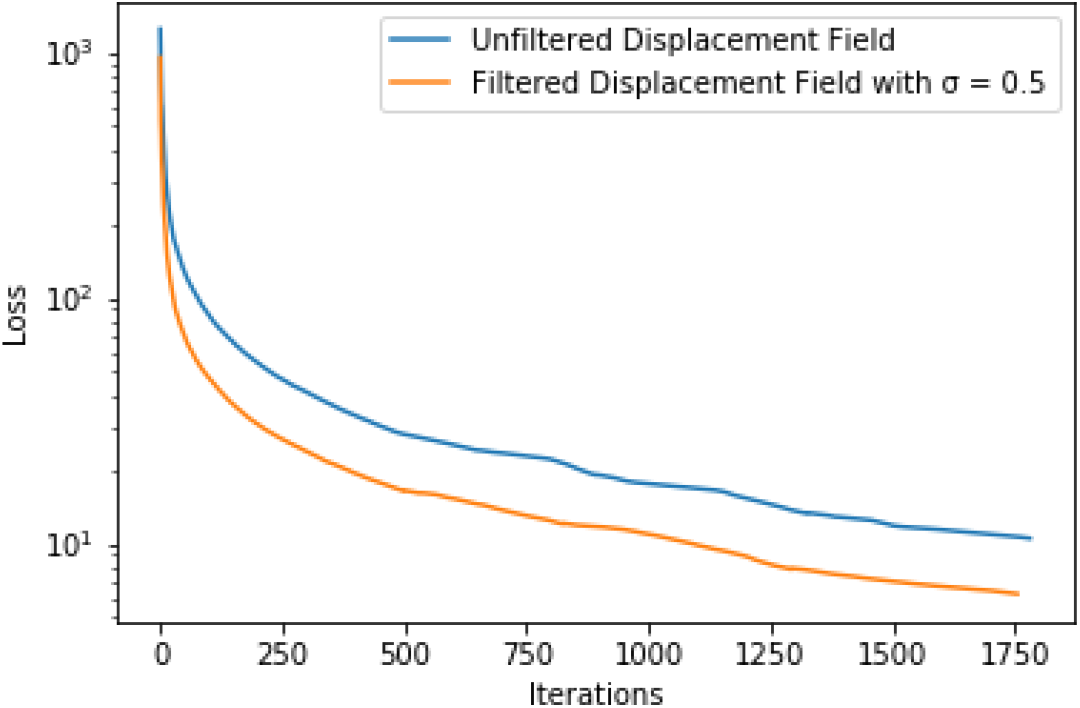
Convergence plot for week 21 to week 22 with and without Gaussian filtering. With the convergence criteria used, we notice that it would require at minimum 1000 more iterations to obtain an additional order of magnitude decrease in the loss.

### 7.4 Results

We obtained the inverse solutions on data without filtering and with Gaussian filtering using zero mean and standard deviation *σ* = 0.5 mm. Table 4 includes results for these cases^2^. Filtering leads to a lower volume-averaged *L*^2^-error between the MRI data and the forward displacement solution driven by the inverted growth deformation gradient field for all cases. However, filtering reduces ∥ ***û***_*τ*_∥_*∞*_ to a degree, especially because surface effects truncate the integrals where the raw displacement is expected to be greatest.

Figure 9 shows the final forward displacement field solution, ***u***_*τ*_, obtained during the adjoint-based gradient optimization using data without filtering. The top and bottom rows in Figure 9, show the inferred displacement field ***u***_*τ*_ on the cortical and ventricular surfaces and are counterparts to Figure 6 which showed the displacement data fields after registration. This comparison provides a visual understanding of how close the inferred 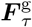 is to the unknown, true 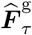, using the displacement fields as surrogates. The second and third rows show the forward displacement fields corresponding to the interpolated ***ũ***_*τ*_ fields. Note that in each case, these are *incremental* fields, for which reason, interpolation into more steps over 23-24 and 24-25 weeks results in smaller magnitudes ***u***_*τ*_. As a result, over 24-25 weeks, in particular, it appears that the forward displacement solution has lower magnitude than the MRI displacement data by registration. The corresponding relative error between the MRI displacement data by registration and the inferred displacement field, ***e***_rel_ = (***û*** *–* ***u***^*h*^)*/*|***û***|, is shown in Figure 10. Higher errors appear in locations of high curvature, e.g. near ventricles and are otherwise homogeneously spread out over the frontal, parietal, and occipital lobes. Table 4 shows, for each stage (by week or at interpolated instants) of the inference, the maximum displacement ∥ ***û*** ∥_*∞*_ and volume-averaged *L*^2^ -norm of the error, as defined in Equation (24). Note that ∥*e*(***u***) ∥_2_ ≤ 2 × 10^*–*2^ ∥ ***û*** ∥_*∞*_ at each stage. Furthermore, on summing *e*(***u***) over the eight steps interpolating between weeks 24 and 25 and using the triangle inequality, it follows that the total volume-averaged *L*^2^-error in the forward displacement field from the inference relative to the MRI displacement data is bounded from above by 4.3 × 10^*–*2^.

**Figure 9.**
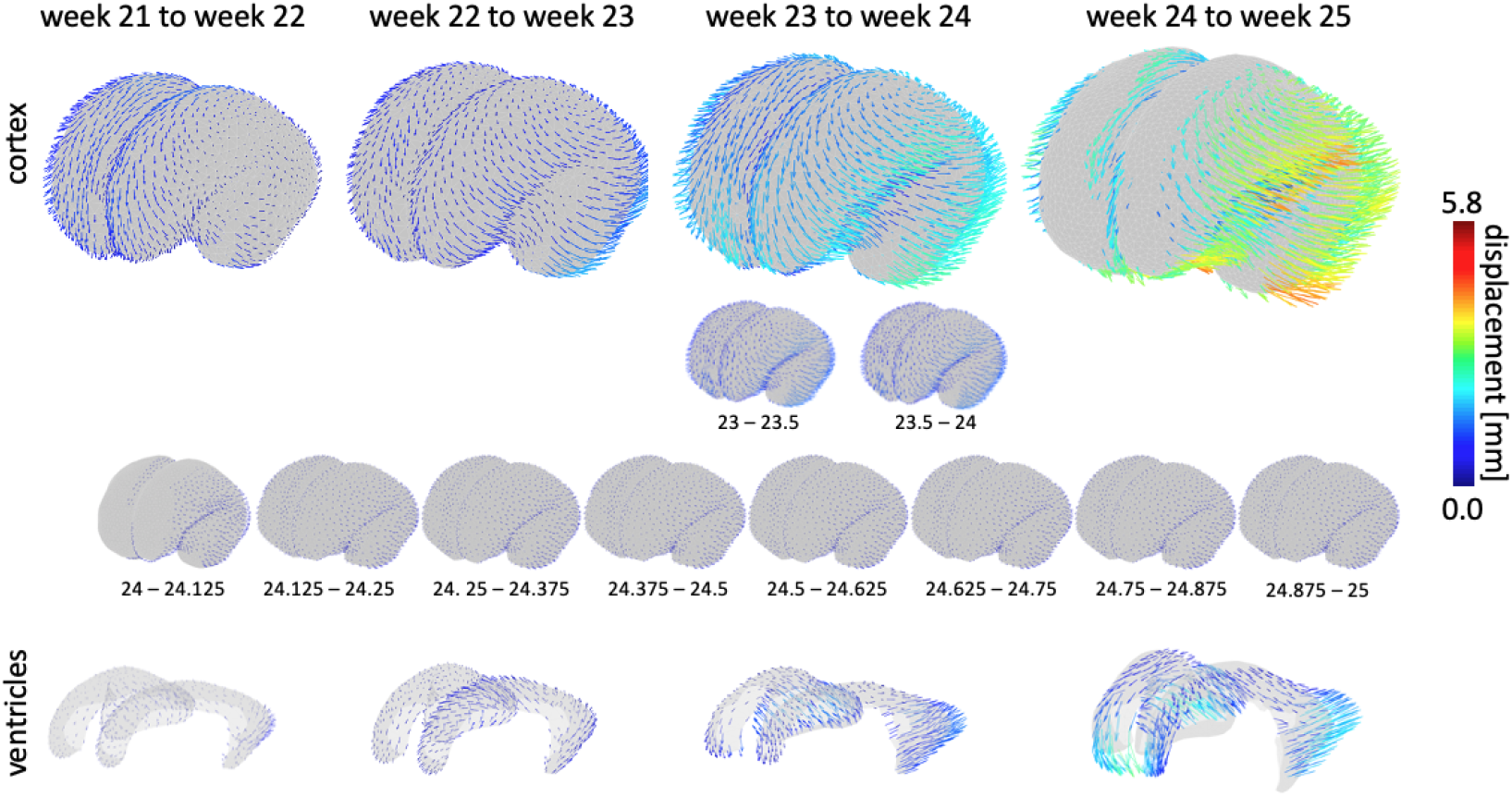
Inferred displacement fields between weeks 21-22, weeks 22-23, weeks 23-24 and weeks 24-25 using our adjoint-based optimization approach and shown here for the cortical and ventricular surface. Changes between weeks 23-24 and weeks 24-25 are broken into 2 and 8 substeps, respectively. Magnitude and orientation of the displacement vectors show remarkable agreement with the registration results shown in Figure 6.

**Figure 10.**
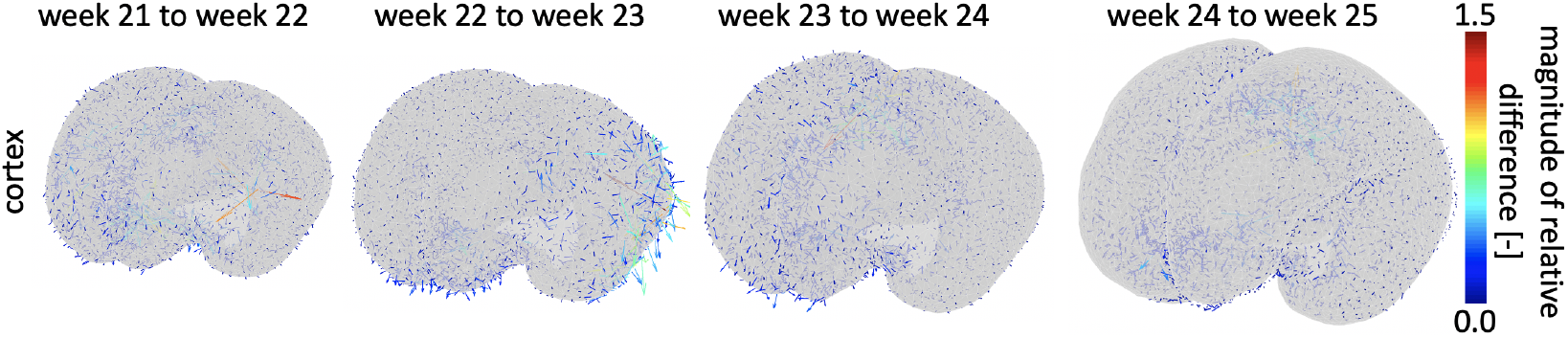
Relative error vectors representing the relative difference between the registration-based displacement fields and the inferred displacement fields. Maximum relative error is smaller than 1.5 and is primarily observed in locations of high curvature, e.g. near ventricles. Generally, we observe a homogeneous distribution of the magnitude and orientation across all weeks. *Arrow size is amplified by factor 10 for visualization purposes*.

The main goal of this study is the inference of 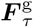 fields at the time instants, *τ*, from adjoint-based gradient optimization. Following inference and before plotting in the figures that follow, these fields were smoothed by Gaussian blurring (Equation 30 with *σ* = 0.5) in order to minimize artifacts introduced by mesh topology. Figure 11 shows the volume change induced by growth alone—i.e., discounting elastic deformation—via det 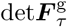 on three representative slices: the coronal, axial and sagittal planes, respectively. Recall that 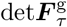 is the volume change induced at each stage *τ* by cell division and growth following migration. Figure 11 therefore offers, to our knowledge, the first data-driven inference of these cell dynamics that are the cause of morphoelastic brain growth, and ultimately of its folding. We draw attention to the radial distribution of growth, seen best in the axial, coronal and sagittal sections and increasing from lower values near the ventricles to higher in the cortex. This distribution is the first data-driven confirmation of the assumption underlying the morphoelastic theory of brain folding: that growth is radially distributed, increasing along the ventricular-cortical direction. The interpolation of morphoelastic growth displacements over eight steps between 24 and 25 weeks, combined with the treatment using evolving reference configurations, renders the inferred 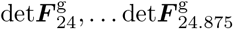 smaller than 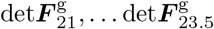. The same pattern of radially distributed growth, increasing from ventricles to the cortical surface, is seen for the eight steps between 24 and 25 weeks when plotted over a narrower range on the right in Figure 11. Also, the volume-averaged det***F***^g^ in the cortex is larger than in the subcortex in all cases, except for growth between 22 and 23 weeks (see Table 5). This strain mismatch between the layers is additional quantitative validation of the kinematic assumption commonly used in morphoelastic growth theories and that drives the emergence of folding, wrinkling and creasing.

**Figure 11.**
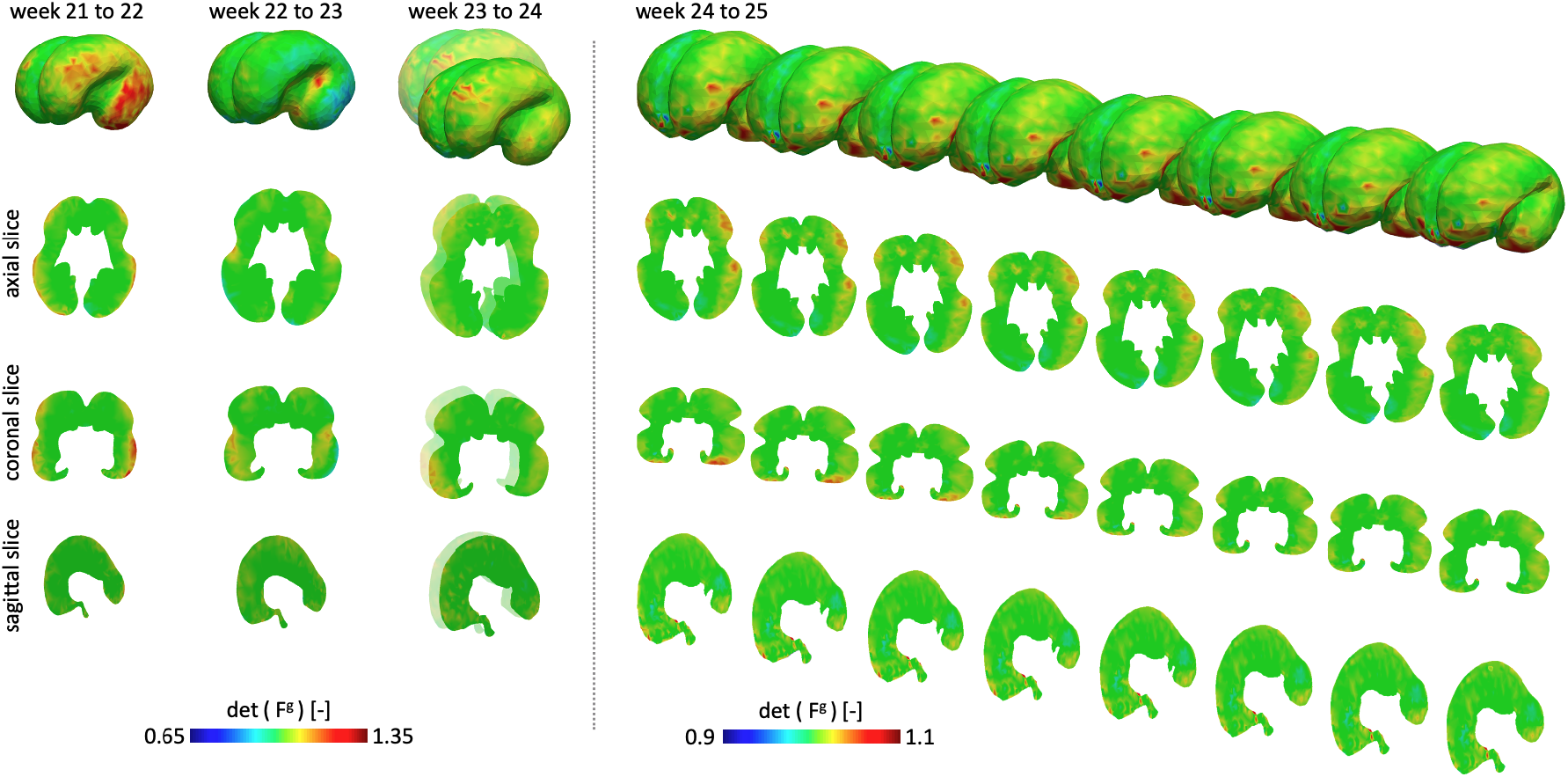
We calculate the determinant of the inferred growth deformation tensor det***F***^g^ for weeks 21-22, 22-23, 23-24, and 24-25 and show them here on the 3D geometry, as well as in representative axial, coronal and sagittal slices. The determinant ranges from 0.65 to 1.35 for weeks 21-24 and range from 0.9 to 1.1 for week 24-25 indicating both localized shrinking and expansion behavior. While growth between weeks 21-24 is mostly homogeneous, a closer look at changes between weeks 24 and 25 reveals localized growth fields in the frontal and temporal lobes. The growth fields are mostly symmetric with respect to both hemispheres, but differ between individual weeks suggesting a characteristic chronological order to brain development throughout gestation.

**Table 5:** The volume-averaged det 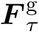, denoted as 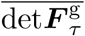 in the cortex is larger than in the sub-cortex in all cases except for weeks 22-23.

| week | $\sigma$ mm | Cortex | | Sub-cortex | |
| --- | --- | --- | --- | --- | --- |
| | | $\overline{\det \mathbf{F}_\tau^g}$ | $\max \det \mathbf{F}_\tau^g$ | $\overline{\det \mathbf{F}_\tau^g}$ | $\max \det \mathbf{F}_\tau^g$ |
| 21-22 | N/A | 1.080 | 1.419 | 1.042 | 1.459 |
|  | 0.5 | 1.058 | 1.239 | 1.036 | 1.371 |
| 22-23 | 0 | 1.031 | 1.413 | 1.041 | 1.398 |
|  | 0.5 | 1.033 | 1.240 | 1.035 | 1.279 |
| 23-23.5 | 0 | 1.071 | 1.321 | 1.048 | 1.337 |
|  | 0.5 | 1.057 | 1.346 | 1.044 | 1.252 |
| 23.5-24 | N/A | 1.071 | 1.367 | 1.048 | 1.364 |
|  | 0.5 | 1.057 | 1.379 | 1.044 | 1.244 |
| 24-24.125 | N/A | 1.021 | 1.190 | 1.013 | 1.140 |
|  | 0.5 | 1.016 | 1.146 | 1.012 | 1.089 |
| 24.125-24.25 | N/A | 1.021 | 1.211 | 1.013 | 1.152 |
|  | 0.5 | 1.016 | 1.165 | 1.012 | 1.096 |
| 24.25-24.375 | 0 | 1.021 | 1.227 | 1.013 | 1.153 |
|  | 0.5 | 1.016 | 1.176 | 1.012 | 1.095 |
| 24.375-24.5 | 0 | 1.021 | 1.229 | 1.013 | 1.136 |
|  | 0.5 | 1.016 | 1.180 | 1.012 | 1.103 |
| 24.5-24.625 | N/A | 1.021 | 1.236 | 1.013 | 1.337 |
|  | 0.5 | 1.0160 | 1.184 | 1.012 | 1.135 |
| 24.625-24.75 | 0 | 1.021 | 1.233 | 1.013 | 1.162 |
|  | 0.5 | 1.016 | 1.185 | 1.012 | 1.166 |
| 24.75-24.875 | N/A | 1.021 | 1.226 | 1.013 | 1.178 |
|  | 0.5 | 1.016 | 1.183 | 1.012 | 1.177 |
| 24.875-25 | N/A | 1.021 | 1.205 | 1.013 | 1.207 |
|  | 0.5 | 1.016 | 1.184 | 1.012 | 1.194 |

While it is suggestive to gain a measure of the total growth over 24 to 25 weeks by multiplying 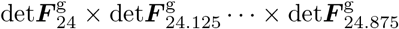, this is not mathematically correct according to the treatment of evolving reference configurations. That is, there is no notion of a quantity, say 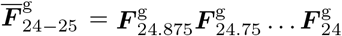 representing pure growth kinematics between 24 and 25 weeks. However, the product of determinants 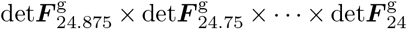 furnishes an approximate value of the total growth-driven volume change. From the consistent appearance of 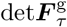 in the frontal and temporal lobes over the eight steps, this approximate measure suggests that the growth volume change ratio between weeks 24 and 25 approaches the range 1.48 to 2.14 in these regions.

Through Figure 11 it also emerges that growth, whose localization is dependent on cell dynamics that is complete by week 26 [41] is indeed focused in the frontal, parietal and occipital lobes, and the cerebellum. Our inverse solutions in Figure 11 indicate that there are some regions where det***F***^g^ *<* 1, implying local contraction dictated by the elastic boundary value problem driven by det***F***^g^ *>* 1 elsewhere.

Figure 12 shows the normal and maximum tangential components of the growth tensor plotted on the corresponding reference configurations for each incremental step. We observe that tangential growth is significantly larger than normal growth which is in line with cortical expansion during early growth followed by cortical thickening during later stages. These results bear comparison with the results of Rajagopalan and co-workers who separately reported the scalar volume changes [33] and principal growth directions [34] resulting directly from the displacement rather than constrained by the laws of morphoelastic growth as we have presented here. This finding of greater growth in the local tangential plane of the cortex than in the perpendicular direction *under the constraint of morphoelastic growth* is the first data-driven confirmation of this mechanism to our knowledge.

**Figure 12.**
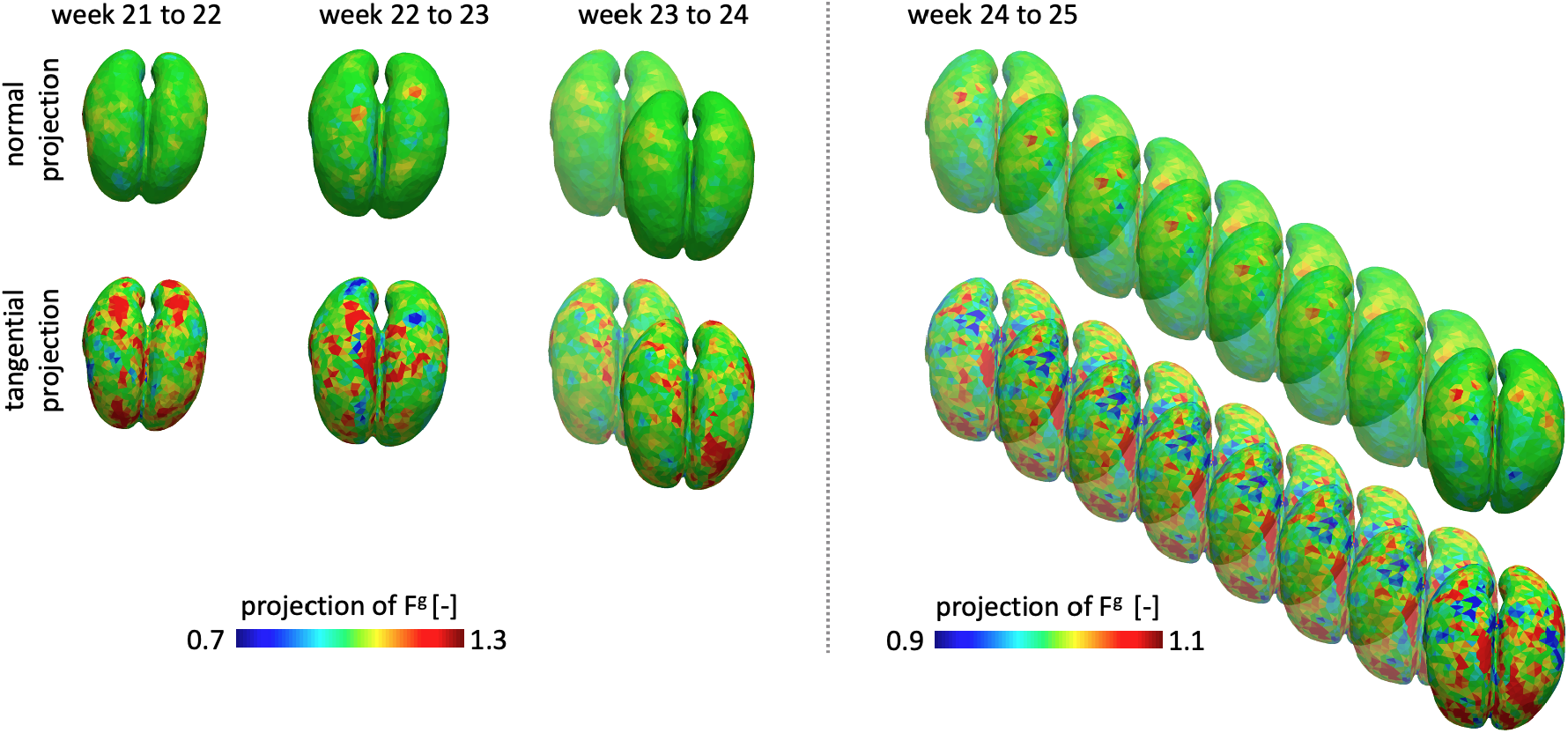
We project the growth tensor ***F***^g^ onto two distinct directions associated with the surface of the brain: the surface normal and a tangent vector. The tangent vector used here provides the maximum tangential projection of ***F***^g^ onto the brain surface. We show both projections for each incremental step of our inversion process and observe that normal growth is much more homogeneous than tangential growth. Strikingly, tangential growth turns out to be much larger than normal growth which supports the notion of in-plane cortical expansion rather than cortical thickening, especially during early fetal brain development.

Finally, we applied Gaussian filtering with zero means and standard deviations *σ* = 1 mm on ***û***_24_. The effect on the inferred 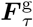 fields from Gaussian filtering of the MRI displacement data (introduced at the beginning of this section) is presented in the Appendix as Figure 13. Larger filters smooth out the displacement fields obtained from MRI data, and also contribute to a more uniform distribution of 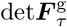.

**Figure 13.**
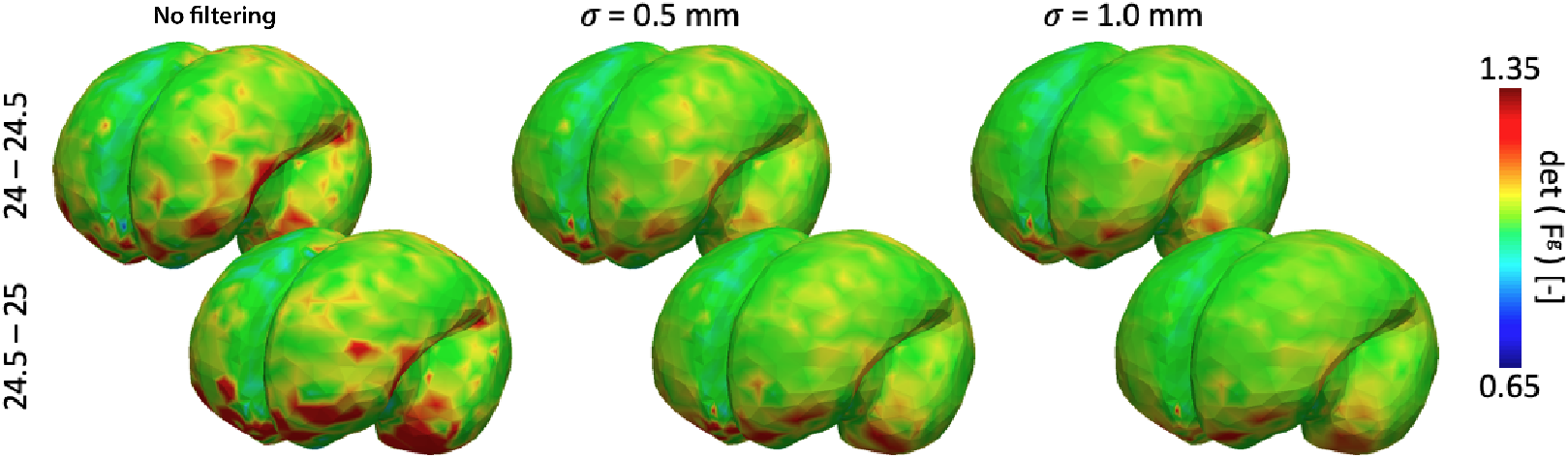
Gaussian filtering with increasing standard deviation *σ* leads to noticeable smoothing of the determinant of the inferred growth deformation tensor det***F***^g^, shown here for the example of changes between week 24 and 25 broken down into two steps.

As explained in Section 7.2, the total deformation gradient tensor 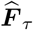 was used as the initial guess for the growth deformation tensor, 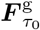. Note that the studies of [33, 34] reported the scalar volume changes and “principal growth directions” from a differently obtained, but essentially the same quantity as 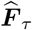. We report the inferred 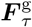 constrained by the laws of morphoelastic growth in Section 7. Nevertheless, it is worth comparing the inferred 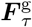 with 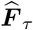, which we present in the Appendix as shown in Figure 14. On comparing with Figure 11 it is clear that det***F*** *<* det***F***^g^, especially in regions of high growth. It follows that det***F***^e^ = det***F*** */*det***F***^g^ *<* 1 in these regions: Local morphoelastic growth leads to elastic compression as indicated by our inverse modelling studies.

**Figure 14.**
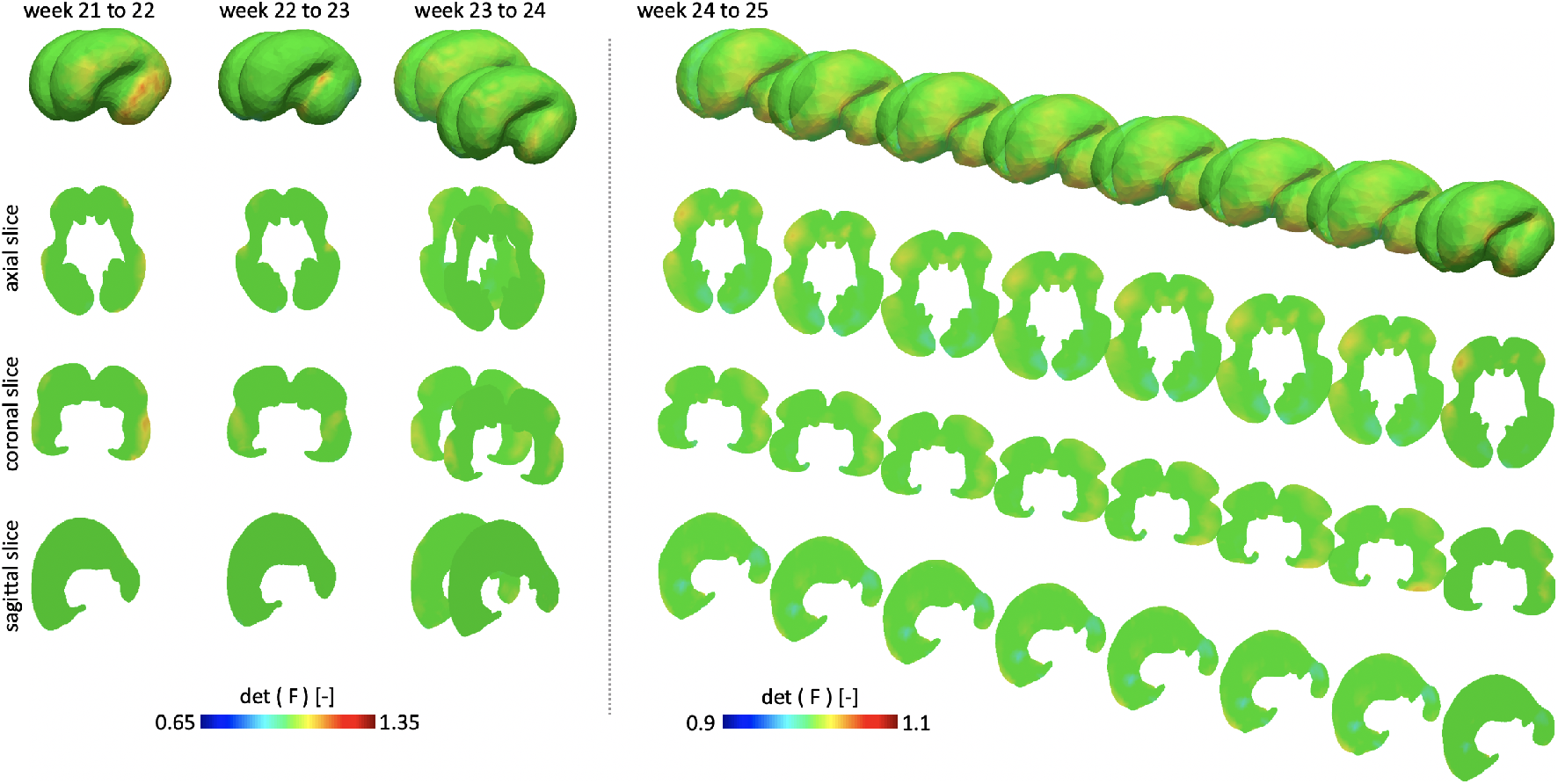
We show the determinant of the total deformation tensor *F* for weeks 21-22, 22-23, 23-24, and 24-25 and show them here on the 3D geometry, as well as inrepresentative axial, coronal and sagittal slices.

## 8 Conclusions

The morphoelastic theory of growth has formed the basis of a large body of computational work on brain development. However, to the best of our knowledge, it has not been used previously to make inferences on the nature of morphogenesis over the course of development. Other data-driven approaches have deduced spatiotemperol variations in the surface growth of fetal brains over weeks 27-37 when the majority of gyrification and sulcification events occur [13]. However, in the aforementioned work, elasticity was used to the extent of minimizing the strain energy for image registration. Our communication takes a step in this direction by building on fetal brain atlases. For it, we have gathered a diversity of methods: MR imaging, segmentation and registration to obtain raw data on the evolving displacement fields that can be regarded as the mapping underlying the geometric changes in the brain over many weeks of development, and inverse modelling to infer the growth tensor via optimization techniques. Notably, the registration techniques that yield displacement data themselves use inverse modelling and *L*^2^ gradient flow-based optimization. The optimization methods that we explored for inferring the growth tensor included gradient descent of a physics-constrained loss function, and separately, adjoint-based gradient optimization, also with the same physics constraint— the satisfaction of the PDEs of morphoelastic growth in weak form. Also notable among our methods is the casting of morphoelastic growth in the framework of evolving reference configurations. Without this version of the morphoelastic growth theory, the problem would become numerically intractable due to the extremely large changes in morphology even over just weeks 21-25 of fetal development.

We note that the results of the inference consistently show that cell dynamics distributes growth radially, increasing from the ventricles to the cortical surface. While the central sulcus begins to form prominently over weeks 24-25, we anticipate that the persistence of this radial distribution leads to the multiscale folding, wrinkling and creasing, whose simulation has been the main goal of previous forward computations of brain morphogenesis—albeit without the strongly data-driven approach that we have adopted here. Our results are broadly in agreement with the premise of Ref. [13] that the growth is larger approaching the cortex, especially in the frontal, parietal and occipital lobes, and in the cerebellum (Figure 11), with some asymmetry between the hemispheres (Figure 12). This qualitative agreement extends to the findings in Ref. [33], which also show greater growth in roughly the same regions of the brain. Figure 12 shows the growth components normal to the cortex and the maximum tangential component in the plane of the cortex. Notably, tangential growth is dominant at this early stage of growth as is broadly understood. The latter findings, while preliminary, are relevant to an understanding of the role of tangential growth and bear comparison with the principal growth direction reported in Ref. [34]. We re-emphasize, however, that unlike the studies of Refs [13, 33, 34], which are based on the observed deformation of the developing brain, ours further subject these deformations to the laws of morphoelastic growth to infer the inelastic, driving growth tensor, ***F***^g^. To our knowledge this is the first confirmation of the dominance of in-plane cortical growth over thickening during early development *under the constraint of the laws of morphoelastic growth*.

The present work serves as a demonstration that the combination of brain atlas data and methods of image segmentation, registration, and finally physics-constrained inverse modelling can provide greater insight to the developmental process. Following this demonstration of a “proof-of-concept”, we intend to carry the study forward through to later stages of development and track the growth tensor as details of primary, secondary, and tertiary gyrification form. Indeed, this is the ultimate goal of the current effort, and will be followed by a stage of linking the macroscopic growth tensor to neuron distribution, the outgrowth of axons and dendrites, and ultimately to pathologies of malformation. We hope to investigate whether, having inferred that growth is concentrated in the frontal, parietal and occipital lobes and the cerebellum, we can extend our techniques to also connecting this growth distribution to the layered distribution of neurons in these cortical regions. This will require high resolution images to resolve smaller fluctuations in growth and other techniques to visualize neuron distributions, and the combination of these data with inference techniques.

In applying these approaches to subsequent stages of development, we anticipate that finer meshes will be needed to resolve the emerging gyri and sulci. The inverse solutions by adjoint-based gradient optimization will require more iterations to attain convergence. Together, these aspects will lead to greater computational expense of our methods. The image registration techniques also will need to be updated in order to delineate morphological features that form between the weekly time instants. These aspects will be addressed in a future communication.

The brain atlas data that forms the basis of our work [18] uses averaging of the geometry obtained from six to eight MRIs at each week of gestation. This approach ensures that only the more repeatable morphological features drive the inference, and also helps with eliminating some of the noise by averaging. However, it does raise the question of whether the resulting geometries satisfy the physics of morphoelastic growth discussed in Section 2. In general, this will not be true, given the nonlinearity of the boundary value problem, and the constitutive response. Having demonstrated the basic viability of our data-driven approaches, we will investigate the extent to which the averaging induces a loss of physical fidelity by also working with data from individual scans in future communications.

## Acknowledgements

This work was supported by the National Institute on Aging of the National Institute of Health under award R21AG067442 to Johannes Weickenmeier, and by the Defense Advanced Research Projects Agency (DARPA) under Agreement No. HR0011199002 to Krishna Garikipati, which also partly funded Zhenlin Wang’s time on this work.

## Appendix

## Footnotes

1 Albeit, solved as elastic unloading from the folded configuration with first-order dynamics added to numerically stabilize the system against bifurcations.

2 Figure 13 in the Appendix shows the det***F*** ^g^ field with an additional level of filtering using *σ* = 1 mm.

